# Lipid scrambling is a general feature of protein insertases

**DOI:** 10.1101/2023.09.01.555937

**Authors:** Dazhi Li, Cristian Rocha-Roa, Matthew A. Schilling, Karin M. Reinisch, Stefano Vanni

**Author notes:** These authors contributed equally (Dazhi Li, Cristian Rocha-Roa, Karin M. Reinisch, Stefano Vanni).

## Abstract

Glycerophospholipids are synthesized primarily in the cytosolic leaflet of the endoplasmic reticulum (ER) membrane and must be equilibrated between bilayer leaflets to allow the ER and membranes derived from it to grow. Lipid equilibration is facilitated by integral membrane proteins called “scramblases”. These proteins feature a hydrophilic groove allowing the polar heads of lipids to traverse the hydrophobic membrane interior, similar to a credit-card moving through a reader. Nevertheless, despite their fundamental role in membrane expansion and dynamics, the identity of most scramblases has remained elusive. Here, combining biochemical reconstitution and molecular dynamics simulations, we show that lipid scrambling is a general feature of protein insertases, integral membrane proteins which insert polypeptide chains into membranes of the ER and organelles disconnected from vesicle trafficking. Our data indicate that lipid scrambling occurs in the same hydrophilic channel through which protein insertion takes place, and that scrambling is abolished in the presence of nascent polypeptide chains. We propose that protein insertases could have a so-far overlooked role in membrane dynamics as scramblases.

## Introduction

A defining feature of eukaryotic cells is the presence of membrane bilayers that separate them from their environment and delineate intracellular organelles with specialized functions. Lipids, the main building blocks of membranes, are primarily synthesized in the cytosolic leaflet of the endoplasmic reticulum (ER) (Vance et al., 1977). From there, since most of them are unable to spontaneously translocate between leaflets due to the associated high energetic cost, they are equilibrated to the ER’s luminal leaflet by integral membrane proteins called “scramblases”, allowing expansion of the ER membrane and vesicles that bud from it (Vance, 2015). Alternatively, lipids from the ER’s cytosolic leaflet can be transported to the cytosolic leaflet of another organelle by lipid transport proteins (Reinisch and Prinz, 2021). At the receiving organelle, lipids must also be scrambled between membrane leaflets to allow for its membrane expansion. This is the case for autophagosomes (Maeda et al., 2020; Matoba et al., 2020; Ghanbarpour et al., 2021), and likely for organelles disconnected from vesicle trafficking pathways, like mitochondria, that rely on protein-mediated transport rather than vesicle trafficking for both their protein and membrane lipid supply (Ghanbarpour et al., 2021; Melia and Reinisch, 2022).

While the role of scramblases in membrane biogenesis and homeostasis is widely accepted (Holthuis et al., 2022; Sakuragi and Nagata, 2023), their identity is mostly unknown, and only a handful of scramblases have been identified and characterized. These include mainly plasma membrane proteins, such as the well-studied TMEM16 (Brunner et al., 2014; Suzuki et al., 2013, 2010) and XK families, that scramble phosphatidylserine during apoptosis (Sakuragi and Nagata, 2023). Lipid scramblases of intracellular organelles have been more elusive (Sakuragi and Nagata, 2023), but recently discovered ER scramblases, VMP1 and TMEM41B, are proposed to work in combination with lipid transport proteins to facilitate lipid transport from the ER (Ghanbarpour et al., 2021; Huang et al., 2021; Li et al., 2021). A common feature of these proteins is the presence of a hydrophilic groove facing the hydrophobic membrane core which allows lipids to slide between hydrophilic membrane surfaces, much like a credit card through a reader (“credit-card model”) (Pomorski and Menon, 2006). Additionally, these proteins may facilitate scrambling by locally thinning the membrane, shortening the distance that lipid headgroups must traverse to cross the bilayer (Falzone et al., 2022). Most likely, one or both of these features are shared by the still unidentified scramblases with roles in membrane biogenesis.

Similar structural features, *i.e.* the presence of a hydrophilic channel in the intermembrane space and the ability to locally thin membranes (Rapoport et al., 2017; Wu and Rapoport, 2021) are shared by another family of integral membrane proteins that is localized in both the ER and organelles disconnected from vesicle trafficking, like mitochondria: protein insertases that translocate peptides across membranes. These structural analogies prompted us to investigate whether insertases could also function as lipid scramblases, thus playing a role not only in non-vesicular protein trafficking but also in non-vesicular lipid transfer. Here we present *in vitro* and *in silico* evidence that lipid scrambling activity is a general feature of protein insertases. We propose that this class of proteins may be among the elusive scramblases with roles in membrane dynamics and expansion.

## Results

### In vitro investigation of insertase lipid scrambling function

To investigate whether insertases have the ability to scramble lipids, we reconstituted a subset of known insertases into liposomes for use in a well-established fluorescence-based lipid scrambling assay (Chang et al., 2004; Kubelt et al., 2002; Menon et al., 2011) (Fig. 1a). In this assay, bovine serum albumin (BSA) is added to liposomes or proteoliposomes comprising a small percentage (0.5%) of short-chain nitrobenzoxadiazole (NBD)-labeled lipids distributed evenly between both bilayer leaflets. BSA extracts NBD-labeled lipids from the outer leaflet of liposomes, and because the fluorescence of NBD-lipids is reduced by ∼50% upon binding by BSA, a ∼25% decrease in fluorescence is observed. In the presence of a scramblase, over time all NBD-lipids in the liposome bilayer become accessible to BSA, allowing for a larger fluorescence reduction of up to 50%, although in practice the reduction is often smaller (35-45%). In contrast to a similar assay that uses dithionite to reduce surface accessible NBD (Menon et al., 2011), the BSA back-extraction assay can also be used with pore-forming proteins (since BSA is too large to enter the liposome lumen through the pore), and it is thus well suited to assay scramblase candidates of unknown structure or oligomeric state, including those whose pore-forming ability is unknown.

**Figure 1.**
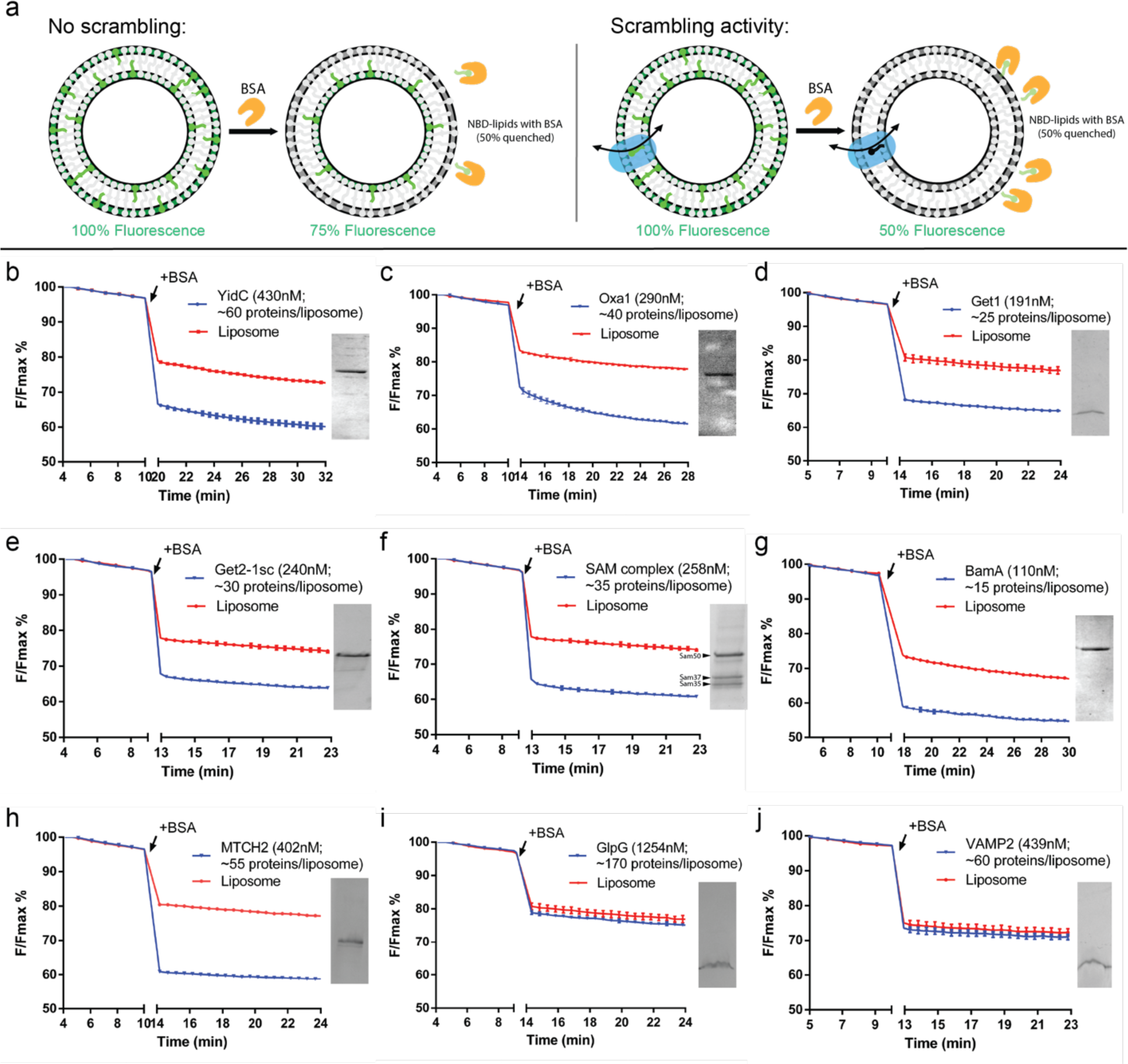
Multiple protein insertases have lipid scrambling activity *in vitro*. (a) Schematic of the BSA back extraction assay. (b-e) Members of the Oxa1 superfamily (YidC, Oxa1, Get1, and the Get1/2 complex) can scramble glycerophospholipids. (f-g) The β-barrel membrane protein insertase, Sam50 in complex with Sam35 and Sam37, and the bacterial ortholog of Sam50, BamA, have scrambling activity. (h) The outer mitochondrial membrane insertase MTCH2 scrambles. (i-j) Negative controls, GlpG and VAMP2, do not scramble. Proteoliposomes used in the assays were analyzed by SDS-PAGE (insets) to confirm efficient reconstitution; approximate numbers for proteins/liposome were estimated assuming 50% recovery of lipids after reconstitution. (See Methods for exact liposome compositions, details of which varied according to experimentalist.)

We reconstituted a recently identified mitochondrial human insertase MTCH2 (Guna et al., 2022) as well as members of the well-studied Oxa1 (Anghel et al., 2017) and Omp85 (Gentle et al., 2004) superfamilies. Both MTCH2 and the Oxa1 proteins feature all-alpha-helical transmembrane (TM) domains, whereas the Omp85 proteins are beta-barrels (Fig. S1). Among Oxa1 proteins, we investigated the inner mitochondrial membrane protein Oxa1 itself, the ER-resident Guided Entry of Tail-anchored proteins (GET) complex (WRB-CAML complex in metazoa), and the *bacterial* insertase YidC. In the Omp85 family, we investigated *bacterial* BamA as well as the Sorting and Assembly Machinery (SAM) complex of the outer mitochondrial membrane. The insertases were isolated in detergent and reconstituted into liposomes using the swelling method (Ploier and Menon, 2016); then the resulting mixture of liposomes and proteoliposomes was further purified by flotation in a density gradient, which allowed removal of unreconstituted proteins and defective liposomes. For those proteins for which the reconstitution into proteoliposomes was less efficient, we also discarded the protein-devoid liposomes in the very top-most fraction of the density gradient. YidC (*E. coli*), Oxa1 (*S. cerevisiae*), the Get1 subunit of the GET complex (*S. cerevisiae*), the GET complex (comprising both Get1 and Get2 from *S. cerevisiae*), BamA (*E.coli*), the SAM complex (comprising Sam50, Sam35 and Sam37 from *S. cerevisiae*), and MTCH2 (*H. sapiens*) all scrambled lipids robustly in the BSA back-extraction assay (Fig. 1b-h). As reported previously, the TM protease GlpG did not scramble (Ghanbarpour et al., 2021), nor did the SNARE VAMP2 (Fig. 1i,j). These data support the hypothesis that scrambling activity might be a general property of insertases.

### High-throughput Coarse-Grain Molecular Dynamics simulations are predictive for scrambling activity by proteins

To more broadly investigate lipid scrambling by insertases, we opted to use molecular dynamics (MD) simulations at the coarse-grain (CG) level of theory, since this methodology has been shown to reproduce well the activity of various scramblases (Jahn et al., 2022; Siggel et al., 2019), and thus provides a cost-effective alternative to experimental approaches. In short, after *in silico* reconstitution of proteins into model lipid bilayers, the inter-leaflet dynamics of all lipid molecules in the system was followed over time, and transbilayer movement of individual lipids was quantified (Fig. 2a).

**Figure 2.**
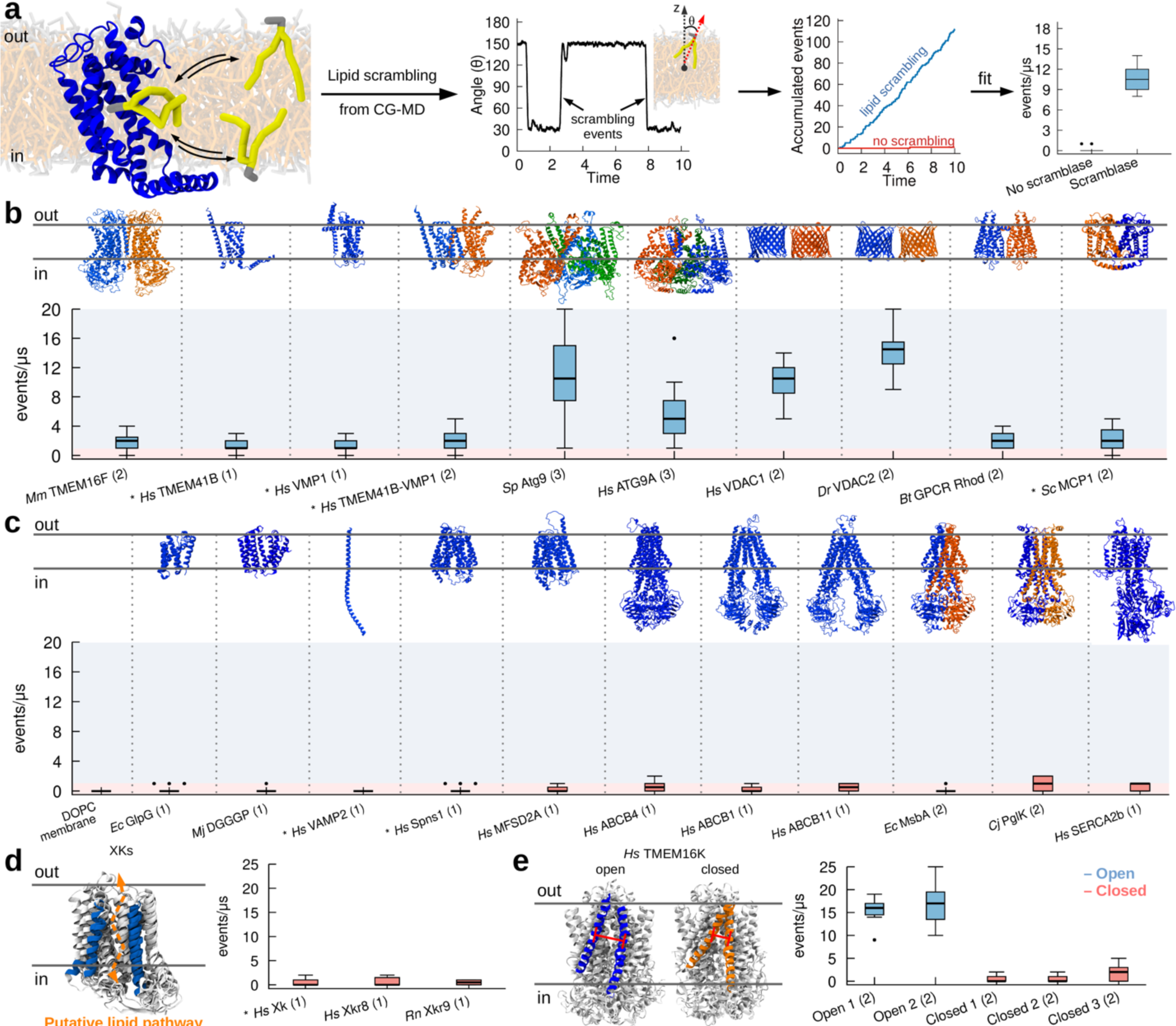
CG simulations recapitulate known activity of lipid scramblases. **a.** Protocol used to quantify lipid scrambling in CG-MD simulations. **b.** CG-MD simulations reproduce lipid scrambling activity by known lipid scramblases of different structure and oligomerization state. **c.** CG-MD simulations correctly reproduce lack of lipid scrambling activity by proteins that do not have scrambling activity *in vitro*. **d,e.** CG-MD simulations recapitulate conformational-dependent lipid scrambling activity by proteins from the XK (**d**) and TMEM16K (**e**) families. AlphaFold structures are denoted by the **\*** symbol, oligomerization state is in parenthesis. Light blue and light red shadings indicate scrambling *vs* non-scrambling activity, respectively. The cut-off used was 1 events/μs.

As a first step, we validated our approach by investigating lipid scrambling *in silico* for multiple known lipid scramblases, ranging over a diverse set of 3D structures, folds, oligomeric states and organisms (Fig. 2b). Our dataset includes TMEM16F (Suzuki et al., 2010), TMEM41B (Ghanbarpour et al., 2021; Li et al., 2021), VMP1(Ghanbarpour et al., 2021; Li et al., 2021), ATG9 (Ghanbarpour et al., 2021; Maeda et al., 2020; Matoba et al., 2020), VDAC1 and VDAC2 (Jahn et al., 2022), Rhodopsin (Menon et al., 2011) and MCP1 (Adlakha et al., 2022) (Fig. 1b). For all proteins, two replicates of 10 μs were run and multiple lipid scrambling events were observed during the MD trajectory, in agreement with the available experimental results. For several proteins, various oligomeric states as reported in the literature were tested (Fig. S2), and the observed trends for lipid scrambling agree with available experimental data. These include, for example, the dimerization requirement for VDAC beta-barrels for proper lipid scrambling, or the higher activity for VDAC2 with respect to VDAC1 (Jahn et al., 2022) (Fig. 2b, S2).

Next, we tested our methodology for several negative controls, *i.e.* proteins that have been shown to not have lipid scrambling activity. In addition to pure lipid bilayers, where no scrambling was observed (Fig. 2c), we investigated three *bona fide* negative controls, the rhomboid protease GlpG (Ghanbarpour et al., 2021), the lipid synthase DGGGp (Ren et al., 2020) and the SNARE protein VAMP2 (this work). In addition, we tested proteins that are not supposed to work as lipid scramblases, such as two lysolipid flippases, Spns1 (He et al., 2022) and Mfsd2a (Chua et al., 2023), and six lipid flippases, ABCB1 (Nosol et al., 2020), ABCB4 (Olsen et al., 2020), ABCB11 (Wang et al., 2020), MsbA (Galazzo et al., 2022), PglK (Perez et al., 2015), and SERCA2b (Yuxia Zhang et al., 2021), including in different conformational states along the lipid flipping cycle (Fig. 2c). In all cases, no or negligible lipid transbilayer movement was observed (Fig. 2c).

Finally, we tested the ability of our MD protocol to discriminate between known active (open) and inactive (closed) states of scramblases (Bushell et al., 2019; Sakuragi et al., 2021; Straub et al., 2021). To this end, we first tested inactive members of the XK family (Sakuragi et al., 2021; Straub et al., 2021) and indeed observed no scrambling in our CG-MD simulations (Fig. 2d). Next we tested different conformations (open *vs* closed) of human TMEM16K (Bushell et al., 2019) and, in agreement with *in vitro* experiments and previous MD simulations (Bushell et al., 2019), we observed scrambling exclusively in the open conformation (Fig. 2e).

Overall, our data suggest that CG-MD simulations can reproduce the experimentally characterized lipid scrambling activity of membrane proteins, including its dependency on protein conformation and oligomerization state.

### Protein insertase complexes have scrambling activity in silico

To further support our hypothesis that protein insertases could function as lipid scramblases, we first used MD simulations to investigate lipid scrambling for several members of the Oxa1 family in their monomeric form. In addition to Get1, Oxa1, and YidC, as in the *in vitro* experiments above, we also investigated MisCB (*B. Subtilis*), OXA1L (*H. sapiens*), Cox18 (*S. Cerevisiae*), Alb3 (*A. Thaliana*), Emc3 (*S. Cerevisiae)*, and TMCO1 (*H. Sapiens)*. Using our approach, we could indeed observe that all tested Oxa1 family insertases can scramble lipids *in silico* (Fig. 3a).

**Figure 3.**
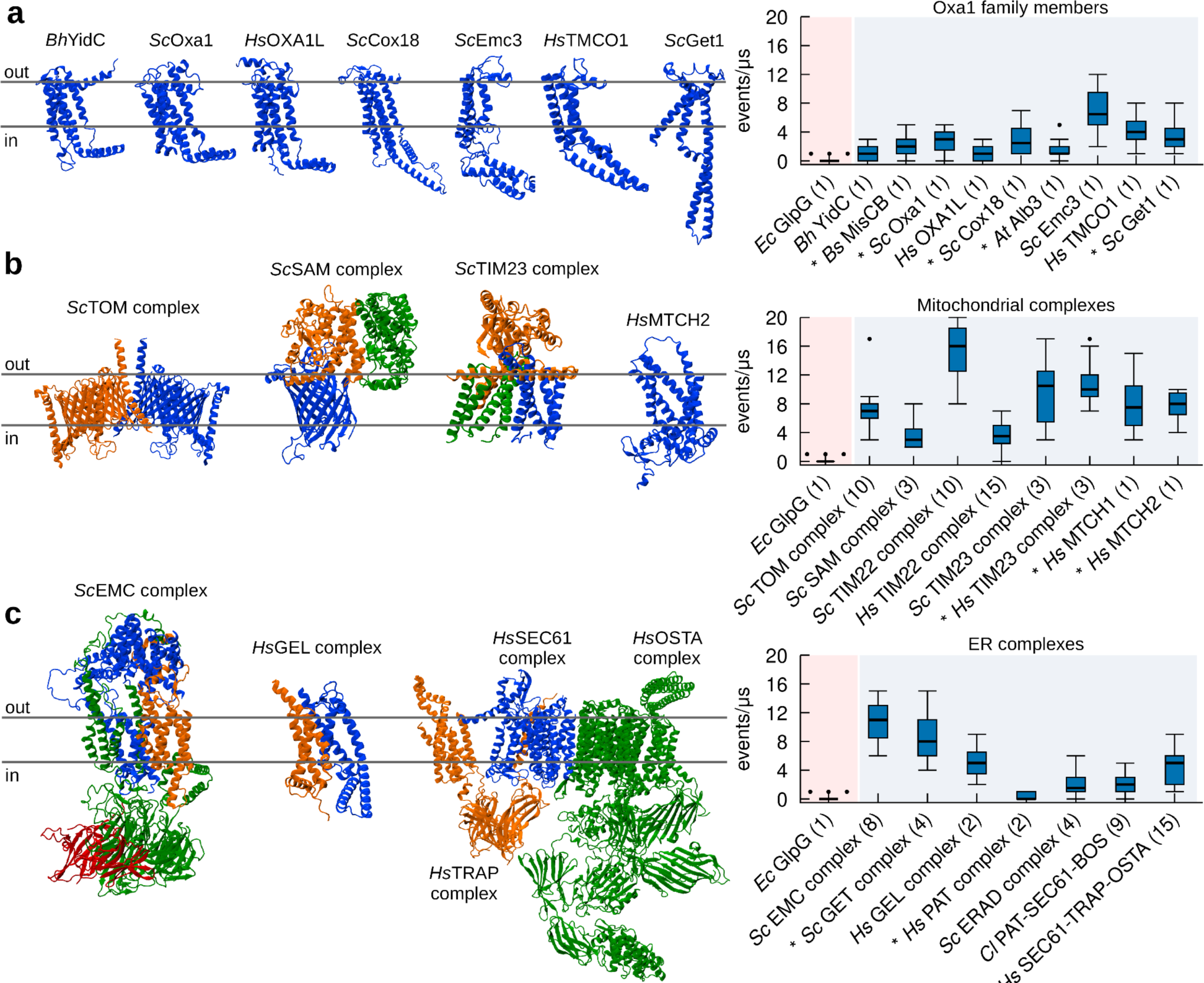
CG-MD simulations identify protein insertase complexes as lipid scramblases. **a.** All members of the Oxa1 family of insertases have *in silico* lipid scrambling activity in their monomeric form. Left: 3D structure of selected members of the Oxa1 family. Right: *In silico* lipid scrambling quantification. The negative control *Ec*GlpG is shown as reference. **b.** Mitochondrial insertase complexes have *in silico* lipid scrambling activity. Left: 3D structure of selected mitochondrial insertase complexes. Right: *In silico* lipid scrambling quantification. **c.** ER insertase complexes have *in silico* lipid scrambling activity. Left: 3D structure of selected ER insertase complexes. Right: *In silico* lipid scrambling quantification. AlphaFold structures are denoted by the **\*** symbol, number of proteins in the complex is in parenthesis.

Next, since several Oxa1 family proteins, such as *Sc*Get1, *Sc*Emc3, and *Hs*TMCO1 are subunits of larger dedicated protein insertion complexes, such as the GET-(McDowell et al., 2020), ER Membrane protein-(EMC) (Bai et al., 2020), and GET- and EMC-like-(GEL) (McGilvray et al., 2020) complexes, respectively, we extended our simulations to all the major protein insertase complexes. In addition to the SAM complex and MTCH2 studied biochemically (see Fig. 1), we investigated the mitochondrial Translocase of the Outer Membrane (TOM) (Tucker and Park, 2019), Translocase of the Inner Membrane 22 (TIM22) (Qi et al., 2021; Yutong Zhang et al., 2021), and Translocase of the Inner Membrane 23 (TIM23) (Sim et al., 2023) complexes, (Fig. 3b). Our results indicate that all these mitochondrial complexes which engage in protein insertion, translocation or assembly into the membrane have clear scramblase activity *in silico* (Fig. 3b). In addition, insertases in the SoLute Carrier (SLC) family such as MTCH1 and MTCH2 also presented scrambling activity (Fig. 3b and Figs S3, S4 and S5).

Next, we focused on the major ER protein insertion complexes. In addition to the GET complex studied *in vitro*, we examined GEL, EMC, Protein Associated with Translocon (PAT) (Sundaram et al., 2022), ER-Associated protein Degradation (ERAD) (Wu et al., 2020), Back Of Sec (BOS) complex (Smalinskaitė et al., 2022), SEC61, TRanslocon-Associated Protein (TRAP) and OligoSaccharylTransferase A (OSTA) complexes (Gemmer et al., 2023) (Fig. 3c). Again, all these ER complexes display lipid scrambling activity *in silico* (Fig. 3c). Interestingly, even for the only insertase complex showing low activity in our simulations (PAT, Fig. 3c) we were able to identify a component with high lipid scrambling activity: Asterix (Fig. S6). A caveat is that in the absence of any experimental structure, we relied entirely on the AlphaFold-derived structures of both Asterix and the complex. In the AlphaFold structure, which is consistent with the cryo-EM structure for PAT in a multipass translocon (Smalinskaitė et al., 2022), Asterix lipid scrambling ability is inhibited by its interaction with its binding partner CCDC47, but we cannot exclude that the AlphaFold prediction for the complex is inaccurate, making a conclusion that Asterix does not scramble in the PAT complex premature.

Notably, our results show that lipid scrambling activity is promoted by specific proteins in the complexes, and that not all components of these complexes scramble lipids (Fig. S3 and S6). Interesting examples in this context are the mitochondrial *Sc*TIM23 complex and the ER *Sc*EMC complex. *Sc*TIM23 is formed by three chains, two of which are integral TM proteins (Tim17 and Tim23) and one exposed to the mitochondrial matrix (Tim44) (Fig. S5). The TM chain reported to be directly involved in protein insertion is Tim17, while Tim23 was suggested not to be involved. In agreement, we observed lipid scrambling exclusively for Tim17 (Fig. S3). One additional component suggested to be a part of *Sc*TIM23, Mgr2, was proposed to act as a seal/cap for Tim17, in relation to the insertion of specific substrates (Sim et al., 2023); notably, when present in our simulations (Fig. S5) Mgr2 reduces lipid scrambling by Tim17 significantly, in agreement with what was previously proposed regarding its role in the *Sc*TIM23 complex. The EMC complex, on the other hand, is composed of eight chains (Emc1-7 and Emc10, Fig. S7), of which at least five are TM (Emc1, Emc3, Emc4, Emc5 and Emc6). Our results are consistent with the fact that both Emc3 and Emc4 are part of the vestibule for protein insertion (Bai et al., 2020), as we observed lipid scrambling only for these two components (Fig. 3a, S6).

Overall, our CG-MD simulations confirm and extend our *in vitro* observation (Fig. 1) that insertase proteins have the ability to scramble lipids. The extent of lipid scrambling depends on protein conformation, oligomerization, and interaction with other members of the insertase complex.

### Lipid scrambling and protein insertion share similar pathways

As all insertase complexes tested have lipid scrambling activity (Figs. 1, 3), we next wondered whether, as hypothesized, lipid scrambling is facilitated by the same hydrophilic groove that promotes membrane protein insertion. Analysis of our trajectories indicate that the main lipid scrambling pathway is localized in the same protein region where protein insertion has been described to take place, and that lipid movement follows a “credit card-like” motion (Fig. 4a). In detail, the mechanism by which lipids are scrambled is mediated by direct interactions between the lipids’ polar heads and protein polar residues in the insertion cavity, thus preventing unfavourable contacts between the polar head of the lipid and the hydrophobic interior of the membrane, and in turn avoiding the interaction between the polar residues located in the insertion region and the hydrophobic body of the membrane. In Oxa1 family proteins, for example, lipid scrambling happens at the hydrophilic interface between 3 highly conserved α-helices in this family (Fig. 4a). Similarly, correlation between insertion and scrambling was preserved in our simulations for *Hs*MTCH2, *Sc*Tim22, and *Sc*Tim17, where scrambling occurs in the described insertion region (Fig. 4a).

**Figure 4.**
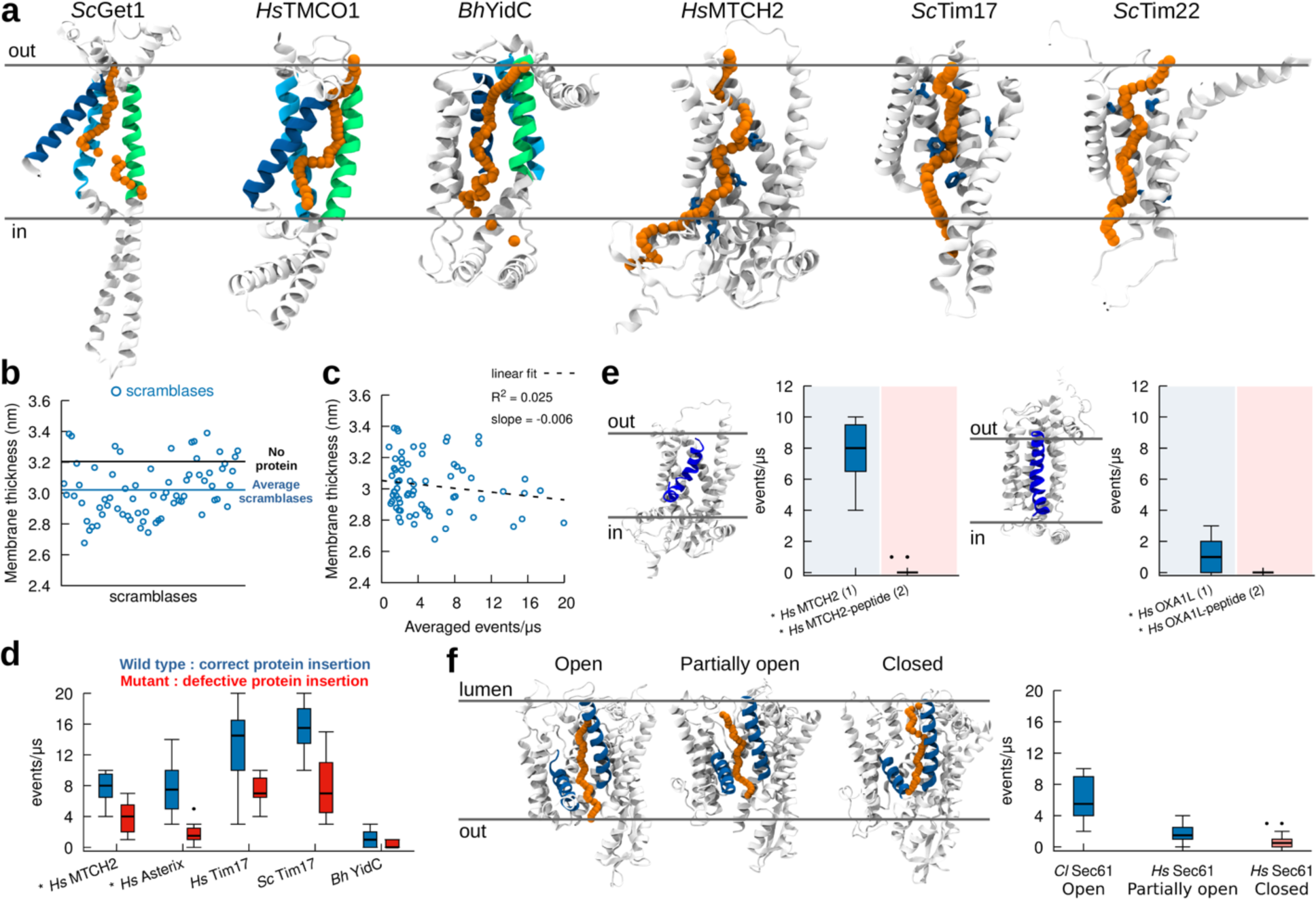
Lipid scrambling takes place via the same mechanistic pathway as in protein insertion. **A.** *In silico* lipid scrambling pathway (orange) in selected protein insertases. The position of the lipid polar head at different times along the scrambling pathway is depicted with orange spheres. Regions involved in protein insertion are shown in green, blue and cyan for Oxa1 family proteins, while residues involved in protein insertion are shown in blue for *Hs*MTCH2, *Sc*Tim17 and *Sc*Tim22. **B.** Protein scramblases induce limited (0.2 nm on average) membrane thinning. **C.** Membrane thickness has minimal correlation with lipid scrambling activity *in silico*. **D.** Mutants proposed to decrease protein insertion activity also reduce lipid scrambling. **E.** The presence of a nascent polypeptide inside the protein hydrophilic cavity abolishes lipid scrambling. Left: *Hs*MCTH2. Right: *Hs*OXA1L. **f.** Different conformations of Sec61 (lateral gate open, partially open and closed) have different lipid scrambling activity. AlphaFold structures are denoted by the **\*** symbol. Number of proteins in the system is in parenthesis.

To further validate this observation, we tried to abrogate lipid scrambling by replacing polar residues in the hydrophilic cavity with hydrophobic ones (Leu), and specifically in the insertase Get1 (Fig. S10). We observed that lipid scrambling activity is very robust, and we could abrogate lipid scrambling *in silico* in Get1 only after 10 mutations were introduced in its hydrophilic channel (Fig. S10). Unfortunately, we were unable to produce and test *in vitro* Get1 with such a high number of mutations as the protein does not fold correctly, and we are thus unable to confirm the importance of the hydrophilic channel via *in vitro* experiments. We did succeed in testing *in vitro* a corresponding mutant (with 10 Leu mutations) for the GET complex (Get1+ Get2) (Fig. S10). However, since Get2 also has partial lipid scrambling ability (Fig. S6), and since the rate determining step in the BSA assay is lipid extraction by BSA rather than scrambling itself, making it possible to detect *in vitro* only complete abrogation of scrambling but not rate reduction, we still observed lipid scrambling for this construct, both *in silico* and *in vitro* (Fig. S10).

In addition to promoting lipid scrambling by favourable interactions with membrane-buried polar and charged residues, we also observed that our dataset of “*in silico*” scramblases also moderately thin (by 0.2 nm on average) the membrane bilayer in its local (R = 1 nm) proximity (Fig. 4b), as previously proposed for lipid scramblases (Falzone et al., 2022). However, we observed only marginal correlation between lipid scrambling and membrane thinning (Fig. 4c), suggesting that while scramblases do indeed thin the membrane, this does not appear to be the main molecular mechanism responsible for lipid scrambling, at least for the dataset of positive lipid scramblases (including protein insertases) we tested.

Overall, our results suggest that lipid scrambling might employ the same mechanistic pathway used in protein membrane insertion. To further test this hypothesis, we performed simulations of insertase mutants that have been shown to reduce protein insertion (*Hs*MTCH2 (Guna et al., 2022), *Hs*Asterix (Smalinskaitė et al., 2022), *Hs*Tim17 and *Sc*Tim17 (Sim et al., 2023), and *Bh*YidC (Kumazaki et al., 2014)). In all cases, we observe reduced scrambling by these protein mutants (Fig. 4d). We further performed MD simulations of two distinct insertases, *Hs*OXA1L and *Hs*MTCH2, in the presence of nascent peptides in the insertion cavity (Guna et al., 2022; Itoh et al., 2021) (Fig. 4e). In both cases, we observe that when the nascent peptide stays in the cavity, no lipid scrambling occurs (Fig. 4e). Finally, for the specific case of Sec61, we tested its ability to scramble lipids in its “closed” and “open” states (Gemmer et al., 2023; Gérard et al., 2020; Itskanov et al., 2023), and our results indicate that when the Sec61 lateral gate is partially closed or closed, lipid scrambling is strongly reduced or abolished, respectively (Fig. 4f). Since the state of the lateral gate has been shown to correlate with protein insertion (Shao, 2023), this result further suggests that lipid scrambling uses the same mechanistic pathway as protein membrane insertion.

## Discussion

Our key finding is that proteins with the capability to insert polypeptide chains into lipid bilayers can also act as lipid scramblases, *i.e.*, they can facilitate lipid translocation from one leaflet to the other. To reach this conclusion, we first used the most reliable approach to investigate lipid scrambling (Ploier and Menon, 2016), biochemical reconstitution of membrane proteins into liposomes together with an *in vitro* scrambling assay. Even though this method is labor-intensive, we succeeded in purifying, reconstituting and assaying 7 insertase proteins/complexes. For all of them, we consistently observe lipid scrambling activity *in vitro*.

Next, using the *in vitro* data, including both our new results as well as previous reports (Suzuki et al., 2010; Ghanbarpour et al., 2021; Li et al., 2021; Ghanbarpour et al., 2021; Maeda et al., 2020; Matoba et al., 2020; Jahn et al., 2022; Menon et al., 2011; Adlakha et al., 2022), as a reference, we established MD simulations as a robust tool for the assessment of lipid scrambling activity. This allowed us to delve more deeply into the molecular mechanisms underlying lipid scrambling, and its correlation with membrane thinning and protein insertion. Moreover, by leveraging the wealth of 3D structures available for insertases and insertase complexes from structural studies (Gemmer et al., 2023; Qi et al., 2021; Sim et al., 2023; Yutong Zhang et al., 2021) and AlphaFold predictions (Jumper et al., 2021; Mirdita et al., 2022), MD simulations allow for large high-throughput screening of proteins and protein complexes in a relatively inexpensive, fast and accurate manner, outcompeting the limited scope of biochemical reconstitution approaches. Specifically, we directly tested *in silico* more than 150 distinct proteins and/or complexes, extending and generalizing our *in vitro* observations.

From a mechanistic perspective, a plausible hypothesis is that protein insertion into the membrane bilayer requires lipid rearrangements, both within and between membrane leaflets. In other words, the scrambling of lipid between leaflets might result in locally-decreased lipid packing, hence lowering barriers for protein insertion. Our observation that lipid scrambling happens in the same groove in which protein insertion takes place suggests that the two mechanisms are unlikely to be simultaneous. Rather, scrambling might precede protein translocation, promoting a poorly packed membrane environment conducive for protein insertion, or follow it to re-equilibrate the membrane bilayer. Overall, even if we were not able to test *in vitro* whether scrambling is required for insertion activity, because peptide insertion activity is more sensitive to mutation than scrambling activity, our results indicate that lipid scrambling could potentially be a general mechanism for local remodeling of membranes. Indeed, in support of our notion that lipid scrambling could alter membrane properties, a recent study indicates lipid scrambling as a mechanism to lower membrane bending stiffness (Wang et al., 2023).

Critically, lipid scrambling can take place independently of protein insertion and, as such, can be considered as a distinct activity. Thus, we propose that protein insertases might have a major, previously overlooked, function in addition to protein insertion, namely as lipid scramblases. Notably, insertases are localized in those organelles that require scrambling activity for membrane maintenance or expansion. These include not only the ER, where most integral membrane proteins enter the secretory pathway and most lipids are synthesized, but also organelles that are disconnected from vesicular trafficking and rely on both protein-mediated protein and lipid delivery, like mitochondria. Our data support a model where a single class of proteins, insertases, play two roles in non-vesicular trafficking: in protein trafficking, their currently recognized function, and also in lipid transfer. Intriguingly, and consistent with this notion, it has been suggested that certain protein insertases might localise at membrane contact sites, where protein-mediated lipid transport also takes place (González Montoro et al., 2018; Koch et al., 2023).

While the molecular mechanisms underlying protein-mediated lipid transport are not yet well understood, an emerging model posits a partnership between bridge-like lipid transfer proteins and scramblases (Ghanbarpour et al., 2021; Melia and Reinisch, 2022). Specifically, for a membrane bilayer to expand, lipids delivered to its cytosolic leaflet by lipid transport proteins must be scrambled between the leaflets of the bilayer. In this context, several bridge-like lipid transport proteins are reported to interact with insertases of the mitochondria: Mdm10 in the ERMES complex (Kornmann et al., 2009), which mediates glycerophospholipid transport between the ER and mitochondria in yeast, interacts with the SAM50 complex (Meisinger et al., 2004), shown here to have scrambling activity. ATG2, a lipid transporter mediating diverse functions including autophagosome biogenesis (Osawa et al., 2019; Tan and Finkel, 2022; Valverde et al., 2019), is reported to interact with the TOM complex (Tang et al., 2019), predicted here to have scrambling activity. And both MTCH2 and TOM, also shown here to have scrambling activity, reportedly associate with the bridge-like lipid transporters VPS13A and VPS13D based on high throughput proteomics (Antonicka et al., 2020; Liu et al., 2018). Scramblases in the ER may also be required to re-equilibrate its membrane bilayer as transport proteins extract lipid cargo from its cytosolic leaflet only. Thus, ERMES interacts with the EMC complex in the ER, which we identified as a scramblase, and the interaction is required for phosphatidylserine transport to mitochondria (Lahiri et al., 2014). Hence, SAM50, TOM, MTCH2, and the EMC could support lipid transport as scramblases. As noted before, the scramblases that participate in lipid transport systems have for the most part not been identified, and our results indicate known insertases in the ER and mitochondria as attractive candidates.

Since there are many insertases in the ER and in mitochondria, each with different substrate preferences for insertion, this implies the existence of multiple scramblases in these membranes. However, our results in no way suggest that insertases are the only scramblases, and it is almost certain that still other classes of proteins (but clearly not all integral membrane proteins as per our *in silico* data and previous studies (Menon et al., 2000)) might also harbor scrambling activity. Scramblases may well be exceptions to the current paradigm of one protein-one function. For example, the TMEM16 proteins and VDAC1/2 in the mitochondria were well-characterized as ion channels (Galietta, 2009; Hartzell et al., 2009; Mannella, 1992) and were subsequently shown also to scramble lipids (Jahn et al., 2022; Suzuki et al., 2010). Hence, the potential lipid scrambling activity of other proteins or classes of proteins will need to be examined on a case-by-case basis. For those proteins with other functions additional to scrambling, whether these functions are simultaneous with scrambling or happen independently as a result of different physiological clues (e.g. protein localization, post-translational modifications, protein-protein interactions, etc.) is a promising future research area.

A current limitation of our study is that we are unable to functionally demonstrate the physiological relevance of insertase scrambling activity. This is because scrambling activity likely is highly redundant, because scramblases may have other roles critical for cell survival such as protein insertion or ion conduction, and because scrambling activity is robust and resistant to mutational ablation. The combination of these aspects makes direct testing of our hypothesis in cells very challenging in practice, if not impossible. We posit, however, that our finding that scrambling activity is a general feature of protein insertases indicates its functional importance. A tantalizing possibility raised by this study is that beyond scrambling lipids in the course of protein insertion, insertases function more broadly in membrane lipid dynamics to participate in membrane growth and expansion. This novel concept would help to rationalize a number of puzzling observations connecting lipid metabolism and transport with protein insertases, including, for example, the role of the EMC complexes in ERMES-dependent lipid transport (Lahiri et al., 2014), mitochondrial morphology defects in the absence of MTCH2 (Labbé et al., 2021), abnormal mitochondrial and lipid droplet morphologies in ERAD-defective brown adipocytes (Zhou et al., 2020); the hypersensitivity to saturated fatty acids of GET complex deletion mutants (Ruggles et al., 2014) or failed thylakoid compartment biogenesis in the absence of the Oxa1 family insertase Alb3 (Sundberg et al., 1997). Most membrane defects arising from insertase dysfunction were previously ascribed primarily to misfolding or mislocalization of the insertases’ protein substrates, but they might also reflect dysfunctional membrane dynamics.

Overall, lipid scramblases have been elusive players in membrane homeostasis, and their identities are only recently starting to emerge (Sakuragi and Nagata, 2023). Our study more than doubles the number of known lipid scramblases, describing tens of new ones. Our results highlight that integral membrane proteins could have additional functions beyond those currently known and suggest protein insertases as new players in membrane dynamics, including in non-vesicular lipid transport. We expect that this new concept will turn out to be particularly helpful not only in the interpretation of numerous existing observations, and especially genetic and physical interactions, but will also open new research directions by blurring the lines between the fields of membrane and protein homeostasis.

## Methods

### Materials

All the lipids, including POPC (Cat. #850457C), POPE (Cat. #850757C), Soy PI (Cat. #840044), NBD-PE (Cat. #810151P), NBD-PC (Cat. #810122), and NBD-PG (Cat. #810161P) were purchased from Avanti Polar Lipids. All detergents were purchased from Anatrace. Bio-Beads^TM^ SM2 Adsorbent Media was purchased from BIO-RAD (Cat. #152-3920). The anti-FLAG M2 resin (Cat. #A2220), EDTA-free Roche cOmplete protease inhibitor cocktail (Cat. # 4693159001), and Optiprep density gradient medium (Cat. # D1556) were purchased from Sigma Aldrich. Fatty acid-free BSA was purchased from AmericanBio. The Expi293™ Expression System Kit was purchased from Thermo Fisher Scientific (Cat. #A14635). The powdered Luria Broth and Terrific Broth were from RPI (Cat. #L24060 and T15100), Teknova, and Fischer Scientific (BP9723).

### Reconstitutions

#### Plasmids

The sequence encoding full-length *S. Cerevisiae* Get1 was cloned into the pET-Duet vector with an N-terminal His_6_-tag. The Get2-1sc-His_6_ construct was a gift from M. Mariappan*. The sequences encoding full-length *E. coli* YidC and BamA (including the N-terminal signal sequence of the latter) were cloned into the pET-29 vector with C-terminal His_6_-tags. Residues 43-402 of *S. Cerevisiae* Oxa1, corresponding to the mature protein, were also cloned into the pET-29 vector with a C-terminal His_6_-tag. The Get2-1sc mutant (T421L/K428L/K433L/W459L/Y461L/S495L/G497L/W501L/N505L/N508L in Get1) sequence was synthesized by Genewiz. Condon-optimized sequences encoding full-length *S. Cerevisiae* SAM50 (N-terminally 3xFlag tagged), SAM35 (no tag), SAM37 (N-terminally Strep tagged), and human MTCH2 (N-terminally 3xFlag tagged) were individually cloned into pCMV-10 vector. The GlpG expression plasmid was a gift from Y. Ha*. The glycerol stock of Rosetta2 cells containing the pTW2-Vamp2-His_6_ plasmid was generously provided by the laboratory of J. Rothman*.

#### Expression and purification of proteins

<u>His_6_-Get1, Get2-1sc-His_6_, and the Get2-1sc mutant</u>: Proteins were expressed and purified as previously described with modifications (Zalisko et al., 2017). The Get2-1sc WT and mutant constructs were transformed to LOBSTR BL21 E. coli cells (Kerafast) and Get1 was expressed in E. coli Ros2(DE3)/pLysS (Novagen). The overnight culture was inoculated into homemade TB medium and cultured at 200 rpm, 37°C until OD_600_ reached 0.6. Protein expression was induced with 0.4 mM IPTG at 17°C for 18 hours for Get2-1sc and 37°C for 18 hours for Get1. Cells were harvested, resuspended in buffer A (500 mM NaCl, 50 mM HEPES, pH 8.0, 10% glycerol, 1 mM TCEP), and lysed by five passes through the Emulsiflex-C5 microfluidizer. The crude lysate was centrifuged at 10,000 g for 20 minutes and the supernatant was further spun at 40,000 rpm for 2 hours in a Ti45 rotor. The membrane pellet was homogenized in buffer A supplemented with 1% Anapoe-C12E9 (Anatrace) or n-Dodecyl-N, N-Dimethylamine-N-Oxide (LDAO, Anatrace), and incubated at 4°C for 2 hours with constant mixing. The suspension was centrifuged at 15,000 rpm for 30 minutes in a JA-20 rotor. The supernatant that contains the extracted proteins was incubated with Ni-NTA resin at 4°C for 30 minutes. The resin was then drained in a gravity column and washed with buffer A supplemented with 20 mM imidazole and 0.02% C12E9 or 0.1% LDAO. The protein was eluted with buffer A supplemented with 250 mM imidazole and 0.02% C12E9 or 0.1% LDAO. The elution was concentrated in a 50K MWCO Amicon concentrator and loaded onto the Superdex 200 10/300 column equilibrated with buffer B (200 mM NaCl, 50 mM HEPES, pH7.4, 5% glycerol, 1 mM TCEP) supplemented with 0.02% C12E9 or 0.1% LDAO (Fig. S11). The peak fractions were pooled and concentrated, and aliquots were frozen and stored at −80°C until use.

#### 3xFlag-MTCH2 and the SAM complex

200 ug constructs encoding 3xFlag-MTCH2 or 1:1:1 mixture of SAM35, Strep-SAM37, and 3xFlag-SAM50 were transfected with Expitransfectamine (Gibco) to 200mL Expi293 cells at a density of 2.5 million cells /ml. Cells were enhanced after 18 hours of transfection and harvested after 48 hours of transfection. The cell pellet was resuspended and homogenized in buffer B supplemented with 2 mM MgCl_2_ and 1x protease inhibitor (Roche). The protein was extracted by incubating with 1% GDN (Anatrace) on a rotator for 2 hours at 4C. The cell suspension was centrifuged at 15,000 rpm in a JA20 rotor for 30 minutes, and the supernatant was incubated with Flag resin for 2 hours at 4C. The resin was washed twice with 10mL of buffer B supplemented with 0.02% GDN and incubated overnight with buffer B supplemented with 1 mM ATP and 2 mM MgCl2 and 0.02% GDN. The resin was further washed 2 times and eluted with buffer B supplemented with 0.02% GDN and 0.2mg/ml Flag peptide 5 times with 20 minutes of incubation in between each elution step. The eluted protein was then loaded onto the Superdex 200 10/300 column that was equilibrated with buffer B supplemented with 0.02% GDN (Figure S11). Peak fractions were pooled and concentrated, and aliquots were frozen and stored at −80°C until use.

#### His_6_-GlpG

The construct was transformed into C43 E coli cells. Protein expression was induced with 0.2 mM IPTG when OD_600_ reaches 0.9, and the cells were cultured at 22°C for 18 hours. Proteins were purified in buffer A as described for Get1, except that n-Dodecyl-β-D-Maltopyranoside (DDM, Anatrace) was used for protein extraction and throughout the purification process. The protein was buffer-exchanged into buffer B supplemented with 0.1% LDAO by loading it onto the Superdex 200 10/300 column (Figure S11).

#### VAMP2-His_6_

Protein expression was induced with 0.5 mM IPTG when OD_600_ reaches 0.8, and the cells were cultured at 37°C for 4 hours. After harvesting, the cells were resuspended in buffer C (25 mM HEPES, pH 7.4, 400 mM KCl, 10% glycerol, 0.2 mM TCEP), supplemented with 1mM PMSF and 4%Triton X-100. The cells were lysed and ultracentrifuged at 35,000 rpm in a Ti45 rotor for 30 minutes. The supernatant was collected and incubated with Ni-NTA resin for 2 hours at 4°C. The resin was then washed with buffer C supplemented with 1% Triton X-100 and 50 mM imidazole, followed by buffer C with 1% n-Octyl-β-D-Glucopyranoside (OG) and 50 mM imidazole. The protein was eluted with buffer C supplemented with 500 mM Imidazole and 1%OG. The eluted protein was loaded onto the Superdex 200 10/300 column equilibrated with buffer B with 0.1% LDAO (Figure S11).

#### YidC-His_6_ and BamA-His_6_

Proteins were expressed and purified as previously described (Ni and Huang, 2015; Serek et al., 2004) with modifications. Both constructs were transformed into C43 *E. coli* cells. The overnight cultures were inoculated into LB medium for YidC or TB medium for BamA, and cultured at 200 rpm, 37°C, until OD_600_ reached approximately 0.7. Protein expression was induced at 37°C for 2.5 hours with 0.5 mM IPTG for YidC, and for 4 hours with 1 mM IPTG for BamA. Cells were harvested, resuspended in buffer E (200 mM NaCl, 25 mM HEPES, pH 8.0, 10% glycerol, 0.5 mM TCEP) supplemented with protease inhibitor (Roche); for BamA, protease inhibitors were included at approximately 2x final concentration and lysozyme was also present. Cells were lysed using the Emulsiflex-C5 microfluidizer, the crude lysate was centrifuged at low speed for 30 minutes and the supernatant was further spun at 40,000 rpm for 90 minutes in a Ti45 rotor. The membrane pellet was homogenized in buffer E supplemented with 1% n-Decyl-β-D-Maltopyranoside (DM, Anatrace) for YidC or n-Dodecyl-N, N-Dimethylamine-N-Oxide (LDAO, Anatrace) for BamA, and incubated at 4°C for 2 hours with constant mixing. Additionally, for BamA the membranes were diluted to at least 25 mL per liter initial cell volume prior to solubilization. The solubilized membranes were incubated with Ni-NTA resin at 4°C for at least 30 minutes. The resins were then drained in a gravity column and washed with buffer E supplemented with 20-30 mM imidazole and 0.2% DM or 0.1% LDAO respectively; in some cases, the detergent for BamA was exchanged on the resin to 0.02% DDM and used for all later steps. The proteins were eluted with buffer E supplemented with 300-330 mM imidazole and 0.2% DM or 0.1% LDAO respectively. The elutions were concentrated in 30K (YidC) or 100K (BamA) MWCO Amicon concentrators and loaded onto the Superdex 6 10/300 column equilibrated with buffer E supplemented with 0.2% DM or 0.1% LDAO respectively (Figure S11). The peak fractions were pooled and concentrated with new 100K MWCO Amicon concentrators, and aliquots were frozen and stored at −80°C until use.

#### Oxa1-His_6_

Proteins were expressed and purified as previously described with modifications (Kohler et al., 2009). Both constructs were transformed to BL21(DE3) codon+ E. coli cells. An overnight culture was inoculated in TB medium, which was subsequently cultured at 200 rpm, 37°C until OD_600_ reached approximately 0.7. The cells were placed in a 4°C cold room for approximately 30 minutes, and protein expression was induced at 25°C overnight with 0.5 mM IPTG. Cells were harvested, resuspended in buffer F (500 mM NaCl, 25 mM HEPES, pH 7.0, 12% glycerol, 0.5 mM TCEP) supplemented with protease inhibitor (Roche, 1.66x final concentration) and lysozyme. Cells were lysed using the Emulsiflex-C5 microfluidizer, the crude lysate was centrifuged at low speed for 30 minutes and the supernatant was further spun at 40,000 rpm for 90 minutes in a Ti45 rotor. The membrane pellet was diluted with buffer F to 40 mL per liter initial cell volume, supplemented with 1% n-Dodecyl-β-D-Maltopyranoside (DDM, Anatrace), and incubated at 4°C for 2 hours with constant mixing. The solubilized membranes were incubated with Ni-NTA resin at 4°C for at least 30 minutes. The resin was then drained in a gravity column and washed with buffer F supplemented with 30 mM imidazole and 0.1% DDM. The proteins were eluted with buffer F supplemented with 540 mM imidazole and 0.1% DDM. The elution was concentrated in a 30K MWCO Amicon concentrators and loaded onto the Superdex 200 10/300 column equilibrated with buffer F supplemented with 0.1% DDM (Figure S11). The peak fractions were pooled and concentrated with a new 100K MWCO Amicon concentrator to approximately 40 μM, and aliquots were frozen and stored at −80°C until use. Immediately prior to reconstitution, the thawed 40 μM aliquots were diluted two-fold with buffer F, yielding final concentrations of 20 μM Oxa1 and 0.05% DDM, which were further diluted ten-fold when added to the reconstitution mixture (along with a matched buffer).

#### Liposome preparation

For YidC, BamA, and Oxa1, 90% POPC (w/w%), 9.5% POPE, and 0.5% NBD-PE (or NBD-PG for BamA) were solubilized in chloroform dried under a nitrogen stream, and further dried under vacuum for at least one hour. For all other proteins, 88% POPC (w/w%), 9.5% POPE, 2% Soy PI, and 0.5% NBD-lipid (NBD-PE or NBD-PC) were solubilized in chloroform, dried under a nitrogen stream, and further dried under vacuum overnight. The resulting lipid films were rehydrated in buffer D (200 mM NaCl, 25 mM HEPES, pH 7.5-8, 1 mM TCEP) to generate a 10.5 mM lipid stock. The mixture was incubated at 37°C for 60 minutes with intermittent vortexing every 20 minutes. The sample was subjected to seven freeze-thaw cycles and extruded 31 times against a 200 nm polycarbonate filter in the Avanti Mini-Extruder.

#### Proteoliposome preparation

As described previously (Ploier and Menon, 2016), for a standard 250 μL reaction, 125 μL of the extruded liposomes were mixed with the reconstitution buffer and Triton X-100 to a final volume of 225 μL. The concentration of Triton X-100 was determined by the swelling assay, typically ranging around 3.4-4.5 mM. After 1-2 hours of destabilization, 25 μL of purified proteins at normalized concentrations were added to the mixture and incubated on a rotator for additional 1-2 hours at room temperature. To remove detergents, pre-washed Bio-Beads were added stepwise: the sample was mixed with an aliquot of Bio-Beads (22-28 mg) at room temperature for 1 hour, followed by replacement with a new aliquot (22-28 mg) and mixing for 2 hours. Finally, the sample was transferred to a new tube containing fresh Bio-Beads (44-56 mg) and rotated at 4°C for 16-21 hours. 150 μL of the recovered sample was mixed with 150 μL of 60% Optiprep and layered with 200 μL of 10% Optiprep and 150 μL of the reconstitution buffer in a 0.8 mL tube (Beckman coulter Cat. #344090). The tube was centrifuged at 40,000rpm for 90 minutes in a SW-55 rotor. The floated liposomes were recovered with a final volume of 150 μL and used immediately. For YidC, Oxa1, and BamA, proteoliposomes were selectively harvested from the region directly below the 10%-0% Optiprep interface, which was found to be protein-rich.

#### BSA back extraction assay

The scramblase assay was carried out in 96-well plates at 30°C. In a triplicate setup, 5 μL of either protein-free liposomes or proteoliposomes were added to 95 μL of the reconstitution buffer. NBD fluorescence was measured using the Synergy H1 Hybrid Multi-Mode Reader (BioTek) with excitation/emission wavelengths set to 460/538 nm. To establish the 100% fluorescence baseline, the measurements were taken for 10 minutes until a steady fluorescence signal was achieved. Subsequently, 5 μL of 3mg/mL fatty acid-free BSA was added to each well, mixed thoroughly, and the fluorescence was measured for 10 minutes. Finally, 0.5% Triton X-100 and 5 mM sodium dithionite were added to each well to ensure that the background signal was small enough (∼3%) so that it did not affect the fluorescence readings. For data processing, each fluorescence reading was divided by the corresponding fluorescence baseline value in each sample.

The scramblase assay for Get2-1sc mutant was carried out with FluoroMax+ spectrofluorometer (HORIBA). For each reaction, 50uL of the proteoliposomes were added to 1950uL of the reconstitution buffer. The sample was vigorously stirred and measured for fluorescence at 460/538 nm for 50-70 seconds to establish a stable baseline. 50uL of 1.5mg/mL fatty acid-free BSA was added, and the fluorescence data were collected for another 200s.

### Molecular Simulations

#### 3D structural modelling

All proteins simulated in this work were obtained from either the Protein Data Bank (https://www.rcsb.org/) or AlphaFold (Jumper et al., 2021) predicted models available from Uniprot (https://www.uniprot.org/). For systems containing more than one chain and for which no structure was available, prediction was performed using AlphaFold-Multimer (Evans et al., 2021) which is implemented in ColabFold (Mirdita et al., 2022); for these cases, a total of 24 recycles were used. The various complexes investigated in the text are described below:

#### Mitochondrial complexes

The monomer of the TOM complex consists of five chains (Tom5-Tom6-Tom7-Tom22-Tom40); thus, its dimer (10 chains) consists of two copies of each subunit from the monomer. The SAM complex consists of three chains (Sam50, Sam35 and Sam37, the last two being non-transmembrane proteins).

The yeast TIM22 complex consists of four transmembrane subunits (Tim18, Tim22, Tim54 and Sdh3) and six helical proteins that form a structure like a ring in the intermembrane space (IMS); this structure serves as chaperone to conduct the substrate from the TOM complex to TIM22 complex (Yutong Zhang et al., 2021). Similar is the case of the human TIM22 complex, where fourteen chains form a double ring-like structure in the IMS and only Tim22 is the transmembrane part.

The yeast TIM23 complex is formed by three chains, two transmembranes (Tim17 and Tim23) and one exposed to the mitochondrial matrix (Tim44). One additional component, Mgr2, was suggested to be a part of *Sc*TIM23 and to act as a seal/cap for Tim17 (Sim et al., 2023).

The human TIM23 complex is composed of three chains (Tim17-Tim23 and Tim50) as its yeast homolog.

#### ER Complexes

The EMC complex is composed of eight chains (Emc1-7 and Emc10), of which five chains are transmembrane (Emc1, Emc3, Emc4, Emc5 and Emc6).

The GET complex is composed of two copies of Get1 and two copies of Get2, forming a heterotetramer. The TRC complex, the human homolog of the ScGET complex, is composed of the WRB, CAML, and TRC40 subunits, the human counterparts of yeast Get1, Get2, and Get3, respectively.

The GEL is composed of TMCO1 (member of the oxa1 family) and C20orf24. The PAT complex consists of two subunits, Asterix and CCDC47.

The ERAD complex consists of four chains, Hrd1, Usa1, Der1, and Hrd3, with Hrd1 and Der1 being the major transmembrane components.

The SEC61 complex is composed of its α, β and γ units. The TRAP complex is composed of its α, β, γ, and δ units.

The OSTA complex is composed of RPN1, RPN2, OST4, OST48, DAD1, STT3a, TMEM258 and OSTC. The translocon complex is composed of the SEC1, TRAP and OSTA complex, and its structure was assembled according to (Gemmer et al., 2023).

#### Set-up of peptide-bound OXA1 and MTCH2 simulations

The initial configuration for the MTCH2-peptide structure was based on the work by Guna et al., 2022. The dimeric structure between *Hs*MTCH2 (Uniprot ID: Q9Y6C9) and the transmembrane part (25-residue fragment from Ile118 to Leu145) of one of its substrates tested *in vitro*, the *Hs*OMP25 (Uniprot ID: P57105), was built using AF multimer and ColabFold (Evans et al., 2021; Mirdita et al., 2022).

Similarly, we predicted the dimer structure using AF multimer of the *Hs*OXA1L-peptide structure based on the work of Itoh et al., 2021. This dimeric structure consisted of *Hs*OXA1L (14niport ID: Q15070) and a 32-residue polyalanine peptide. It is worth mentioning that in both cases, the best prediction based on AF corresponded to the peptide located into the well-characterized insertion cavity of each protein.

All systems were subjected to a minimization step in vacuum for a maximum of 50,000 steps or until the maximum force on any atom was less than 100 kJ/mol. For this purpose, the steepest descent algorithm and the AMBER99SB-ILDN force field were used (Lindorff-Larsen et al., 2010).

#### Coarse-grained molecular dynamics (CG-MD) simulations

The minimized structures of each system were embedded in a DOPC membrane using the CHARMM-GUI web server (Jo et al., 2008). Subsequently, CG-MD were carried out using the GROMACS software, version 2019.6 (Abraham et al., 2015), and the Martini 3 force field (Souza et al., 2021). Elastic network was used to preserve the 3D structure of the proteins and the multimers, using a force constant of 500 kJ mol^-1^ nm^-2^. Two replicates of each system were carried out using different initial velocities and for a time of 10 μs, using a time step of 20 fs. The temperature was maintained at 310 K using the V-rescale thermostat (Bussi et al., 2007) and the pressure at 1 bar using the Parrinello-Rahman barostat (Parrinello and Rahman, 1981). Additional information for all simulated systems are shown in Table S1.

#### Calculation of scramblase activity from CG-MD

To calculate scramblase activity, we measured the angle of each lipid with respect to the z-axis. Thus, lipids located in the upper leaflet will have angles ∼0° while lipids located in the bottom leaflet will have angles ∼180° (Fig 2a). We define a buffer region between 55° and 125°, in order to reduce the noise generated by lipid movement and thus not overestimate the events obtained. A scrambling event was counted when a lipid in the upper leaflet moved to an angle greater than 125°, and when a lipid in the bottom leaflet moved to an angle lower than 55°. These angles were calculated using the gmx_gangle tool and were measured every 1 ns. The vectors used to measure these angles were created using C4A-NC3 and C4B-NC3 beads, corresponding to the last beads of the tails and the headgroup bead of a DOPC lipid according to Martini 3 labels. For all analyses the first 2 μs were omitted, thus resulting in 16 data points per simulated system (8 data per replica, each data represents the number of events in 1 μs). These 16 data points were used to build the corresponding boxplots using GNUPLOT.

#### Calculation of the local thickness of the membrane

We extracted the local thickness of the membrane from the curves of the 2D density diagrams for the hydrophobic body of the membrane, *i.e* excluding both the head group and the phosphate group. These 2D densities were calculated with the gmx_density tool along the Z axis, and were calculated every 5 ns omitting the first 2 μs of each trajectory. The densities were calculated considering the lipids that were at a distance of 1.0 nm from the protein (Fig. 3b).

## Acknowledgments

We thank T. Melia, W. Kukulski, and W. Prinz for comments regarding this manuscript, as well as lab members from the Vanni lab and Reinisch lab. We gratefully acknowledge support from NIGMS (R35GM131715 to KMR), the Swiss National Science Foundation (grants CRSII5_189996 to SV), and the European Research Council under the European Union’s Horizon 2020 research and innovation program (grant agreement no. 803952, to SV). This work was supported by grants from the Swiss National Supercomputing Centre under projects ID s1176, s1030 and s1221. DL received support from the China Scholarship Council.

## Supporting Material

**Table S1.**
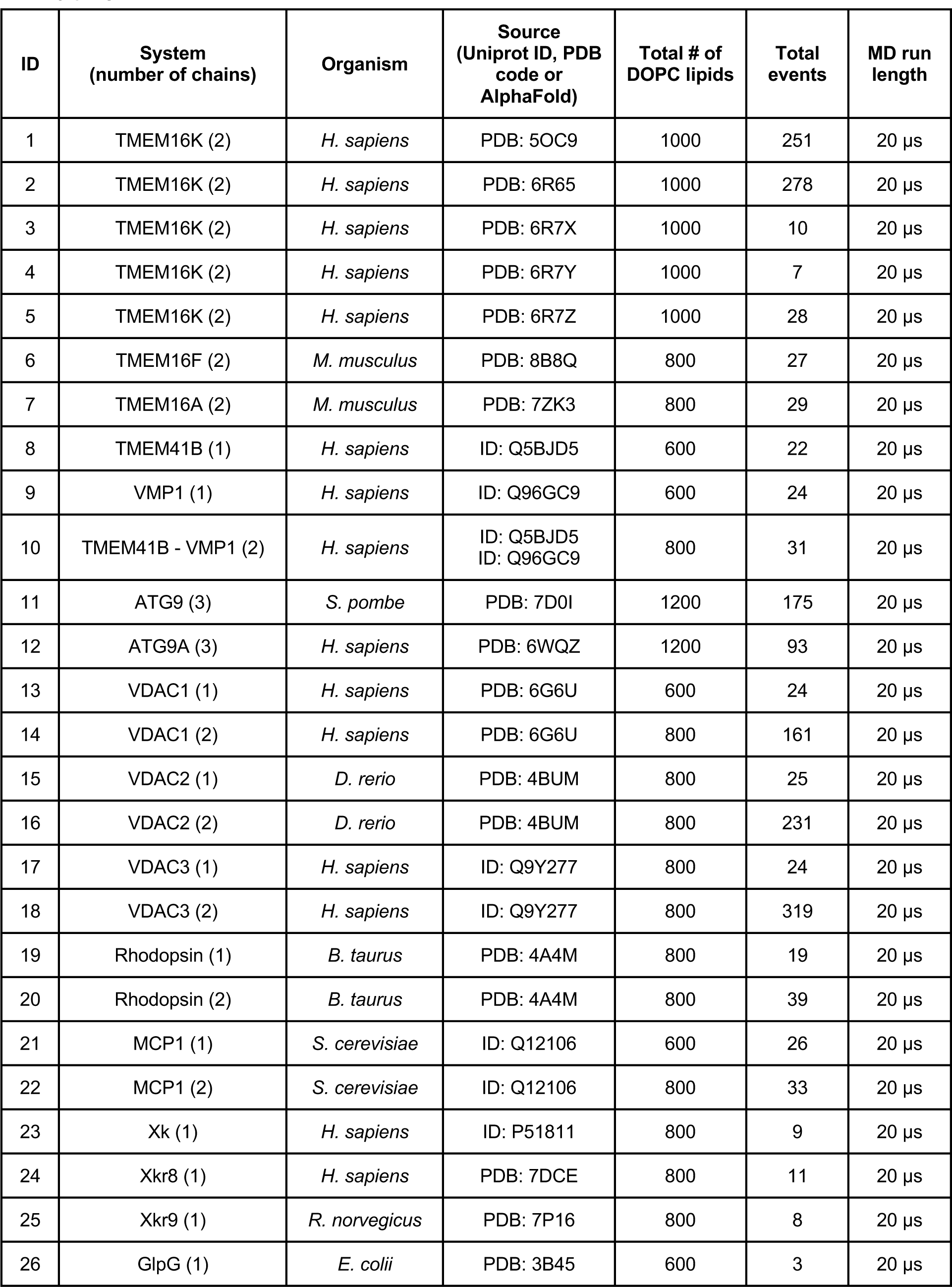
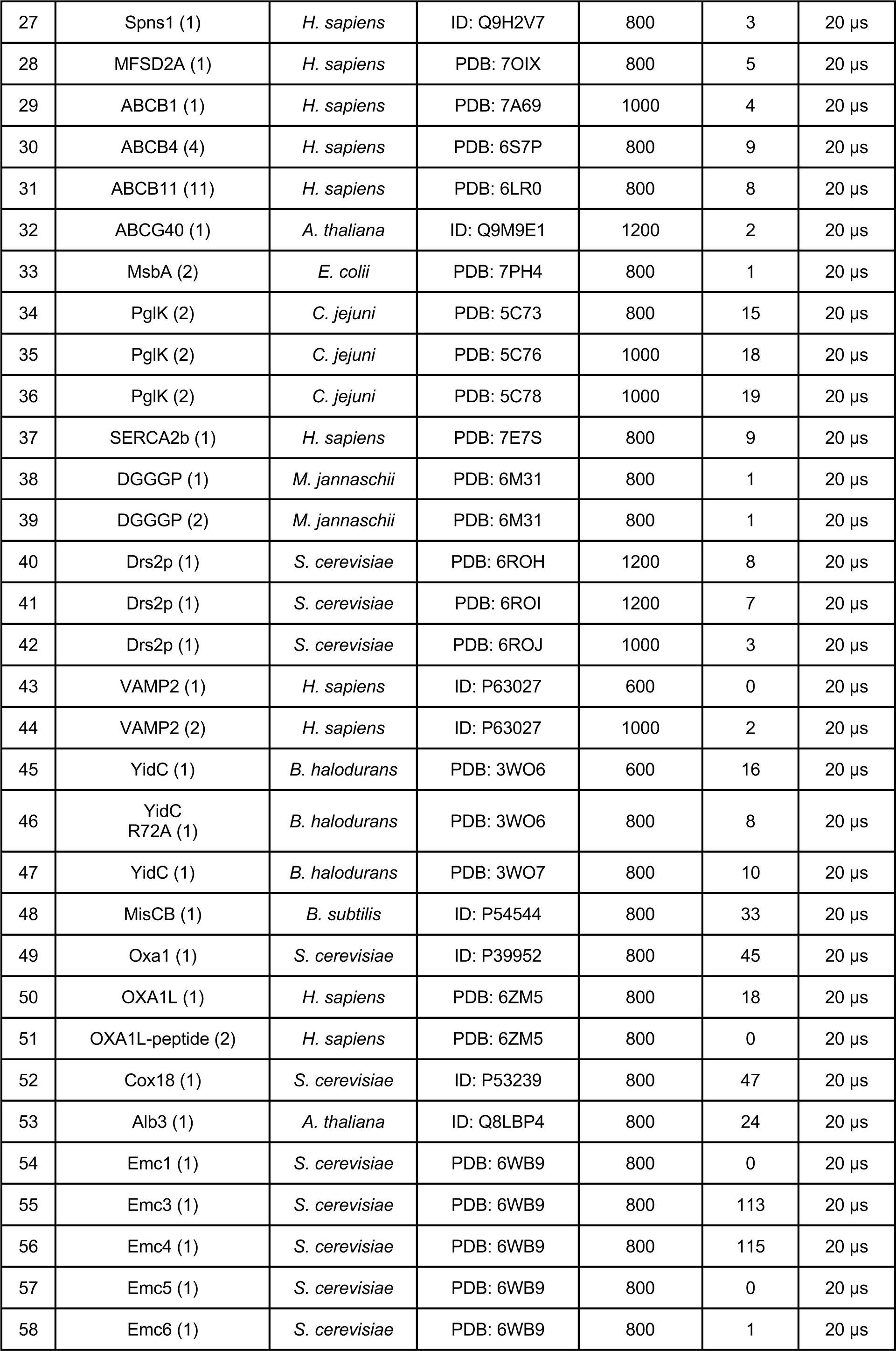
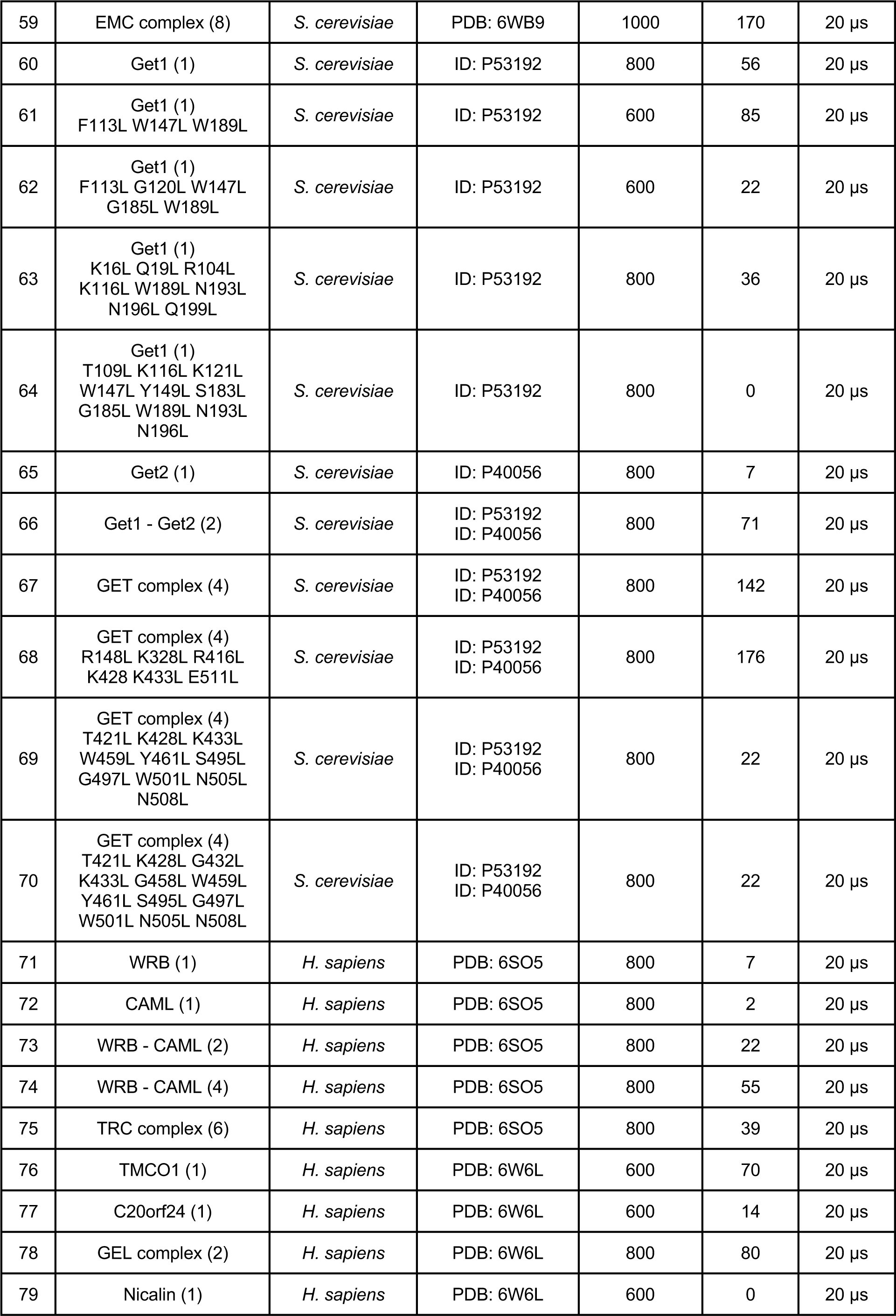
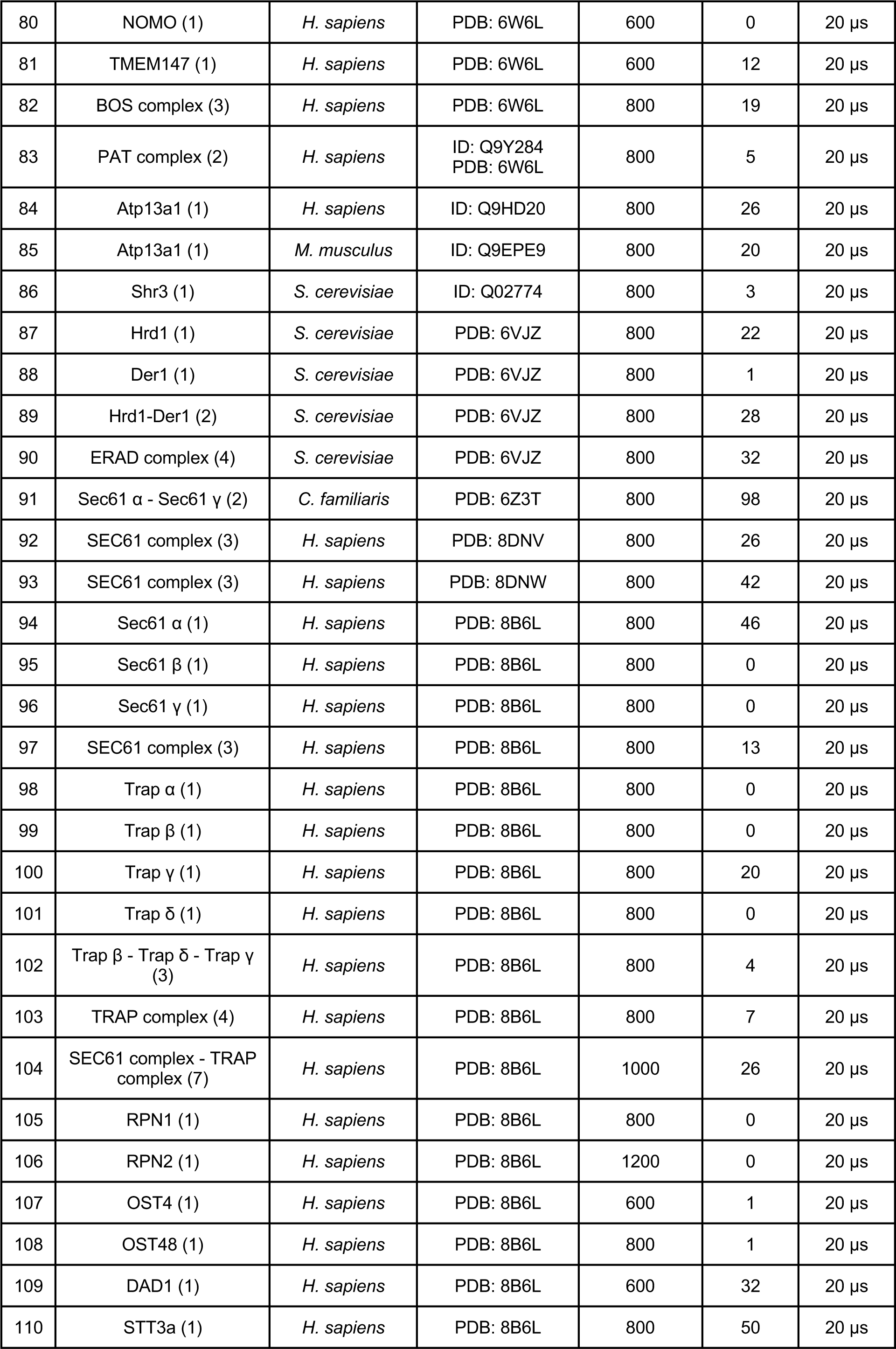
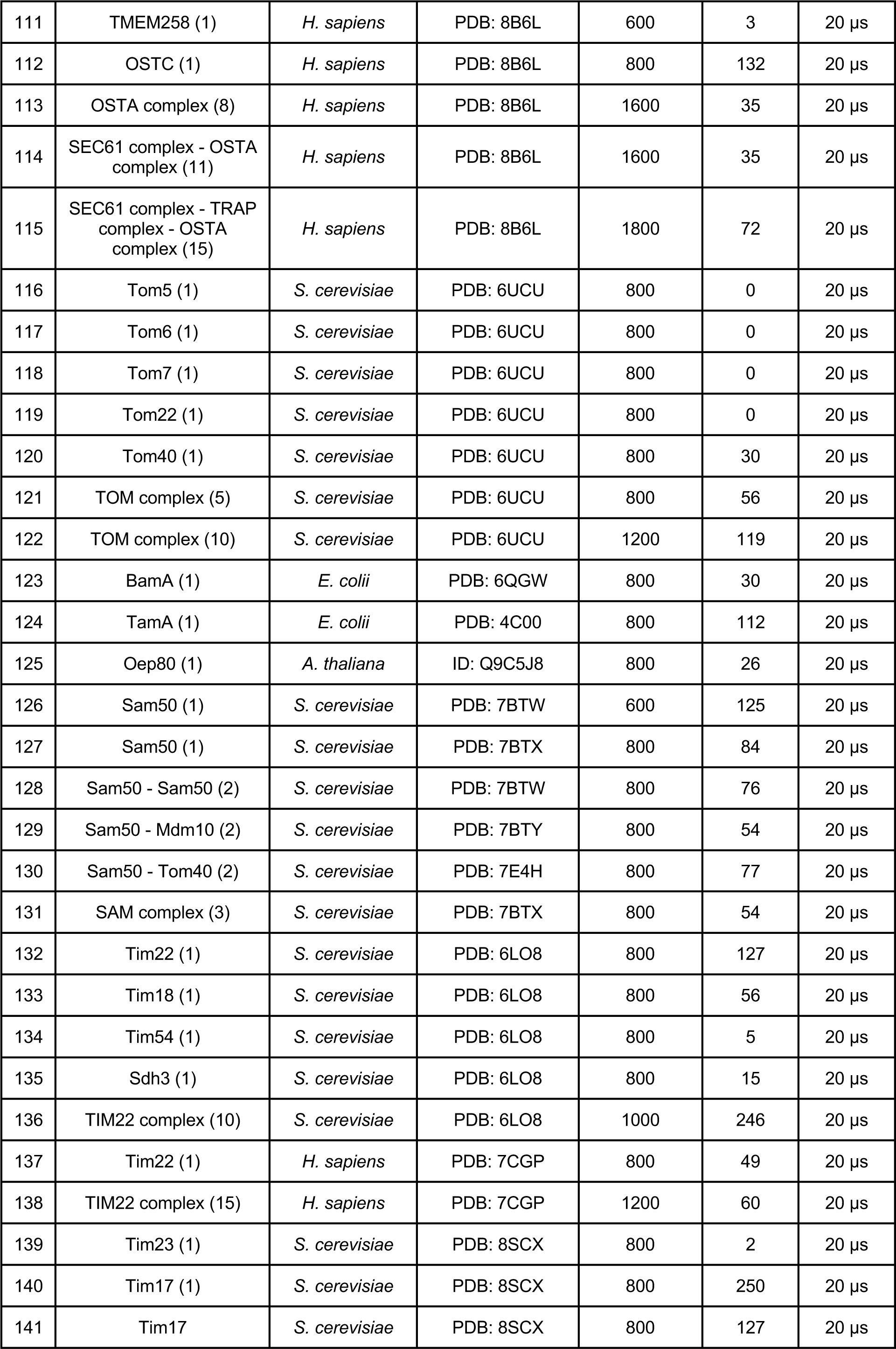
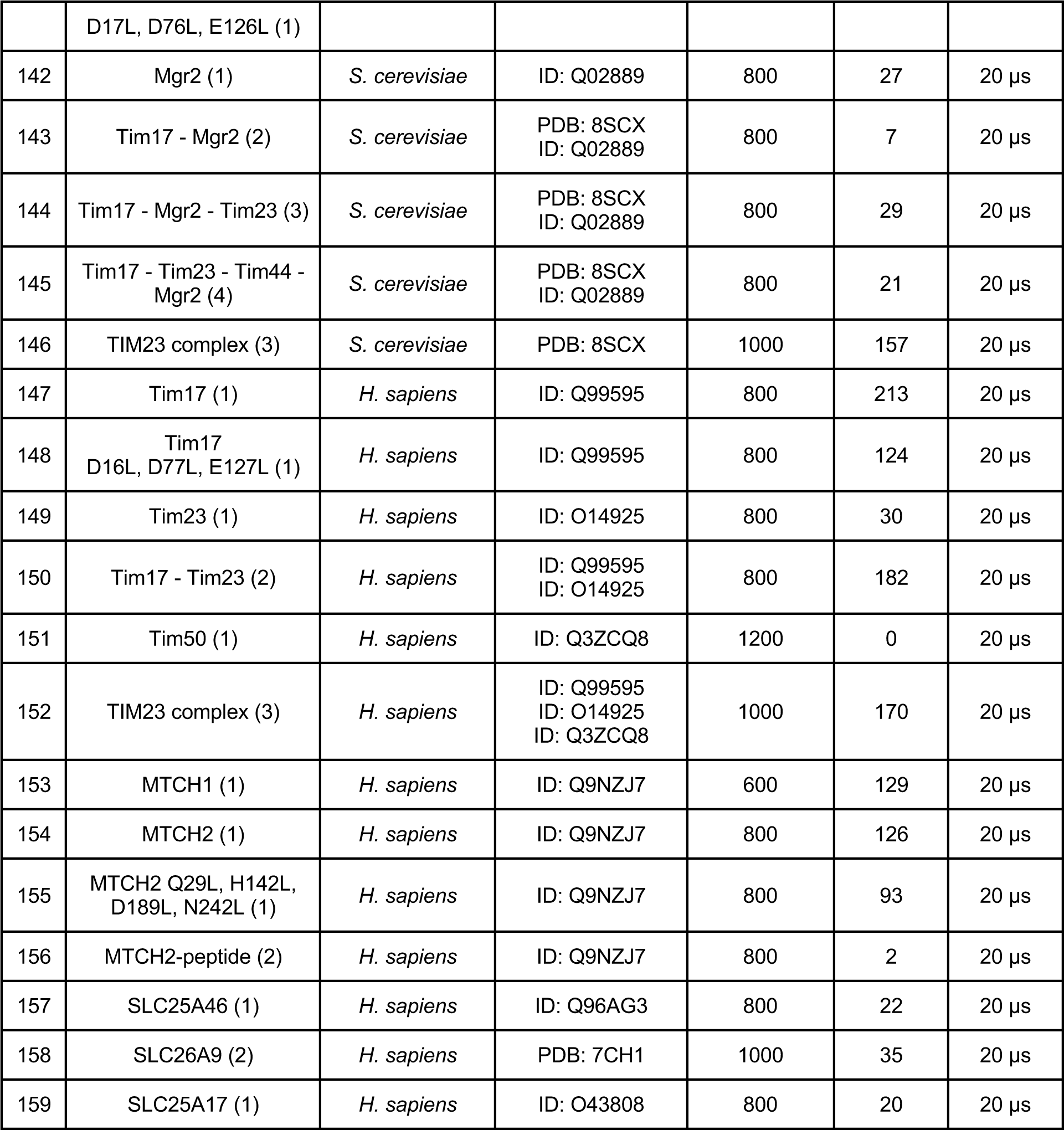
Details about all simulated systems. Systems for which the uniprot ID is provided means that the available AlphaFold (AF) model was used. All oligomers were predicted with AF multimer.

## Supporting Figures

**Figure S1.**
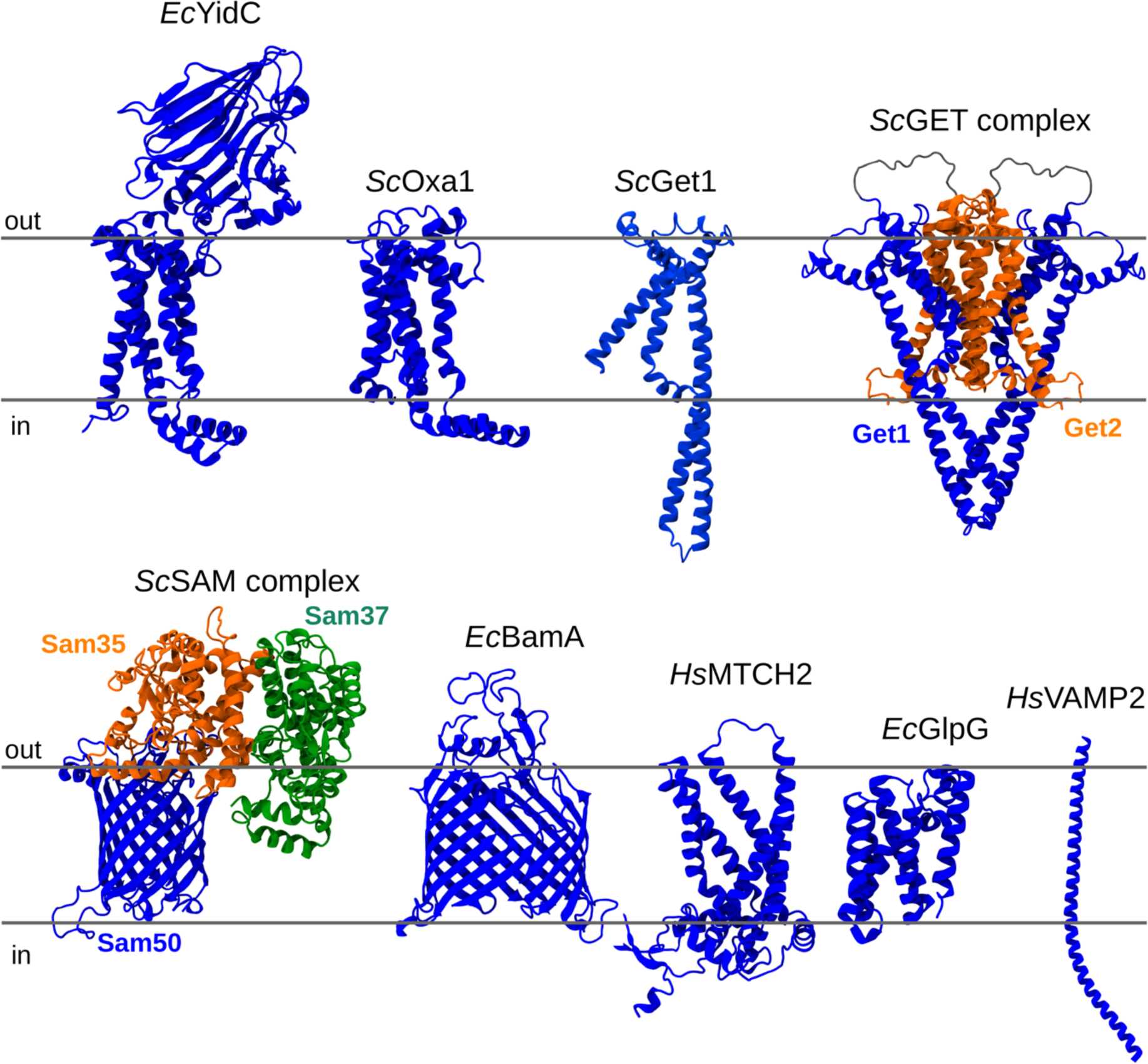
3D structure of experimentally tested proteins.

**Figure S2.**
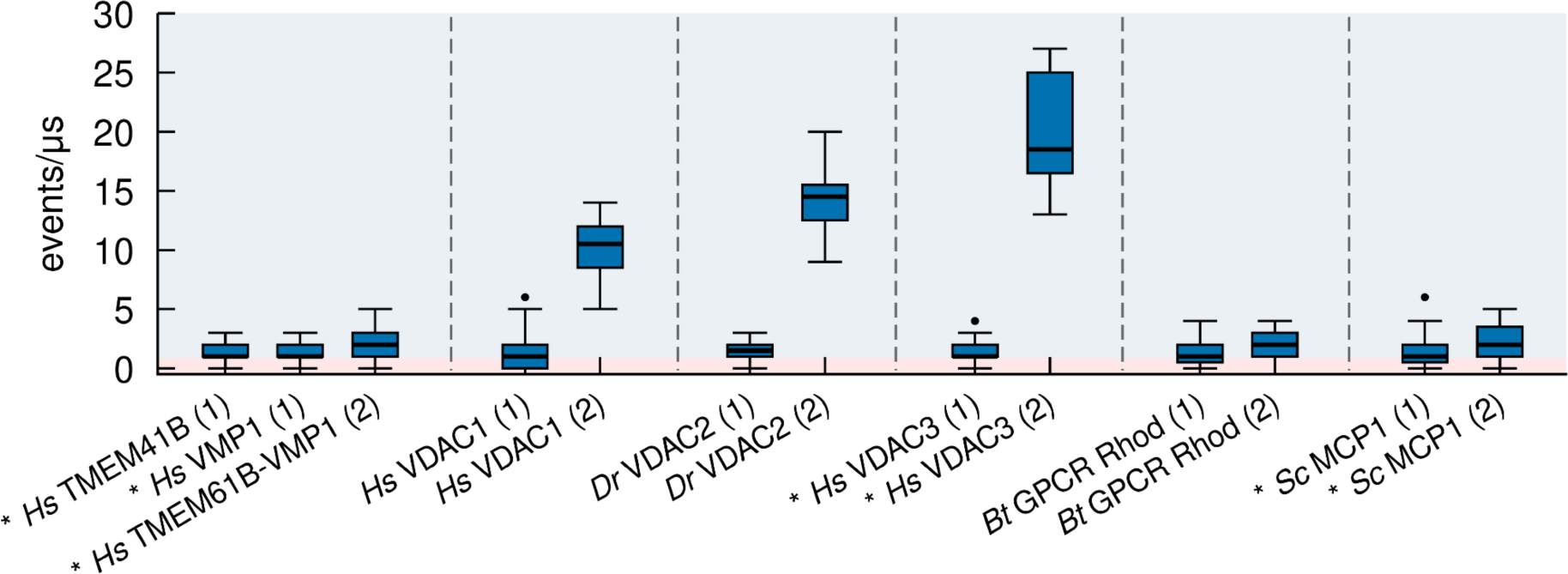
Oligomerization influences lipid scrambling. Comparison of lipid scrambling activity between monomeric and dimeric forms of selected scramblases. The blue shaded region represents the behavior for a scrambling-positive protein and the red area for a scrambling-negative protein. AlphaFold structures are denoted by the **\*** symbol, oligomerization state is in parenthesis.

**Figure S3.**
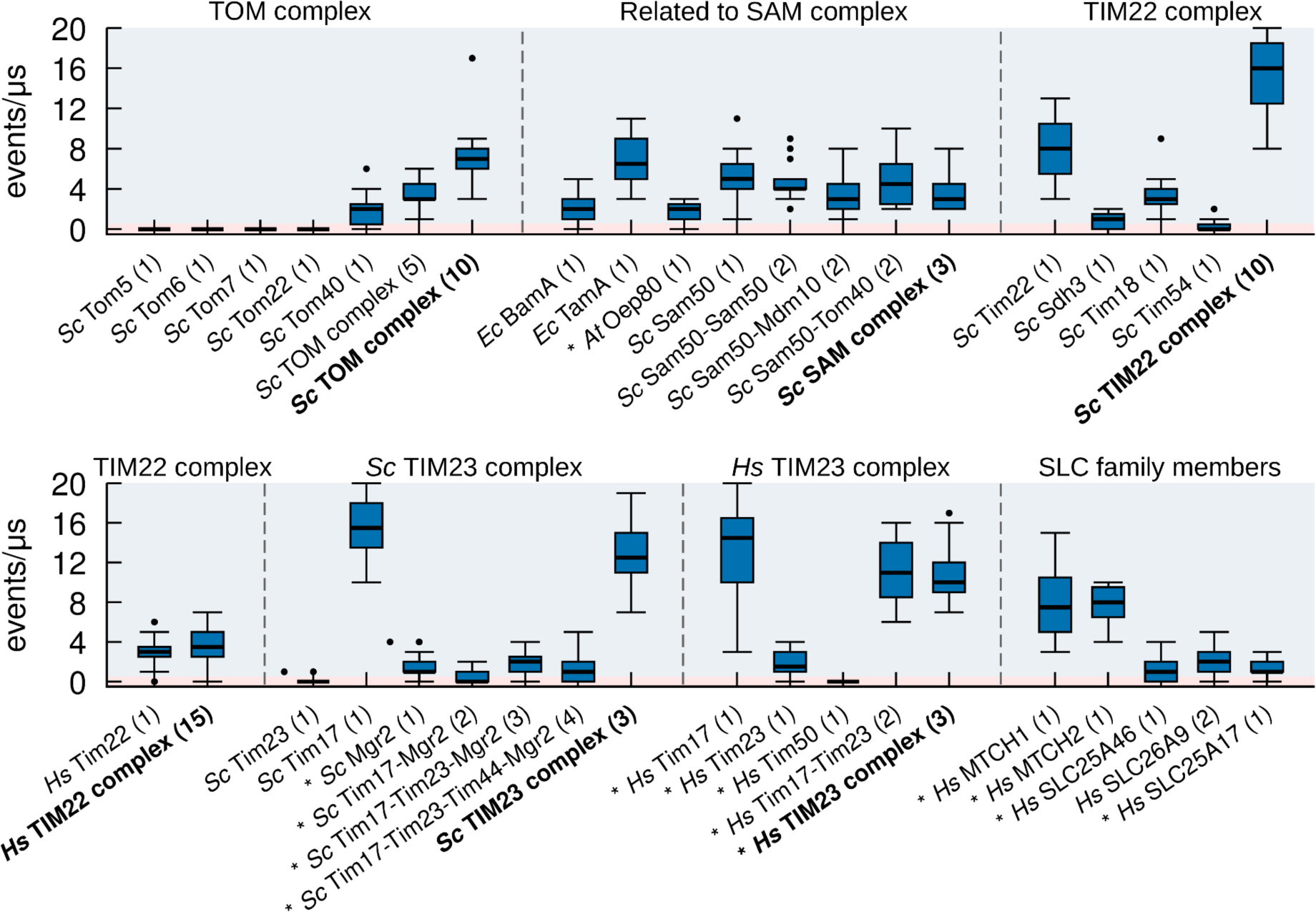
Extended version of insertase complexes in the mitochondria. Lipid scrambling activity of individual components and selected complexes for the major mitochondrial insertases investigated. The blue shaded region represents the behavior for a scrambling-positive protein and the red area for a scrambling-negative protein. AF structures are denoted by the **\*** symbol, oligomerization state is in parenthesis.

**Figure S4.**
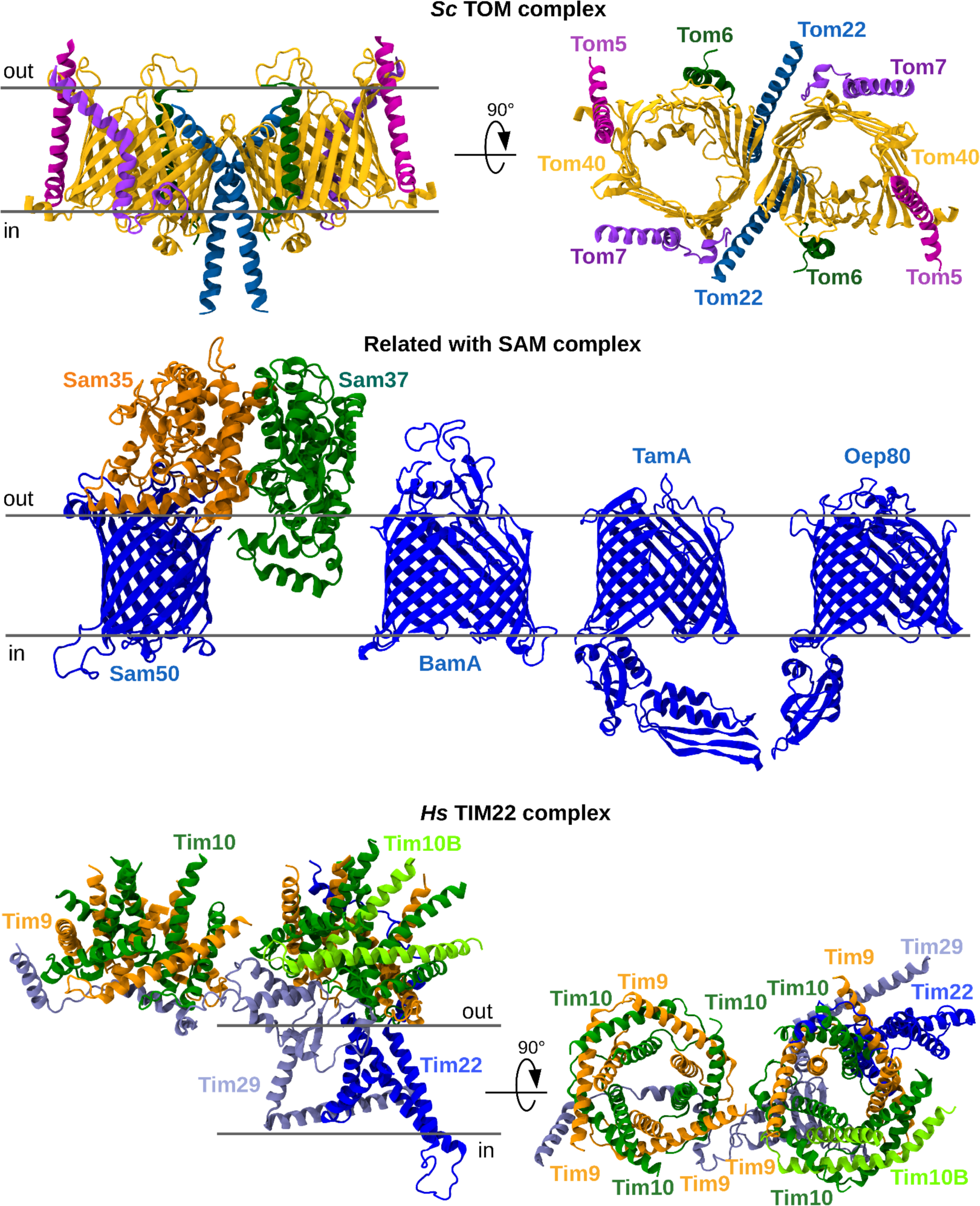
3D structures of mitochondrial complexes TOM, SAM and *Hs*TIM22.

**Figure S5.**
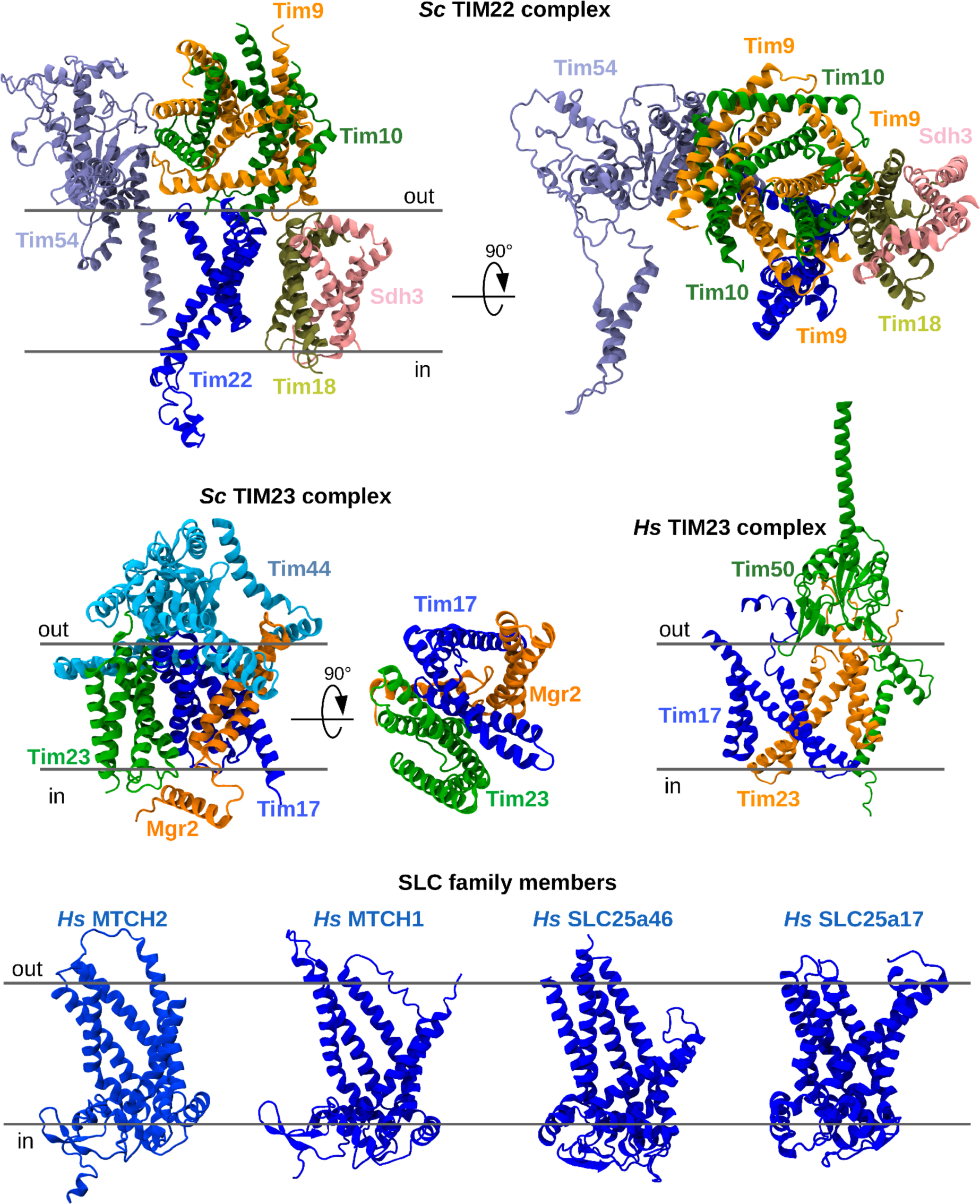
3D structures of mitochondrial complexes *Sc*TIM22, *Sc*TIM23, *Hs*TIM23 and SLC proteins MTCH2, MTCH1, SLC25a46 and SLC25a17.

**Figure S6.**
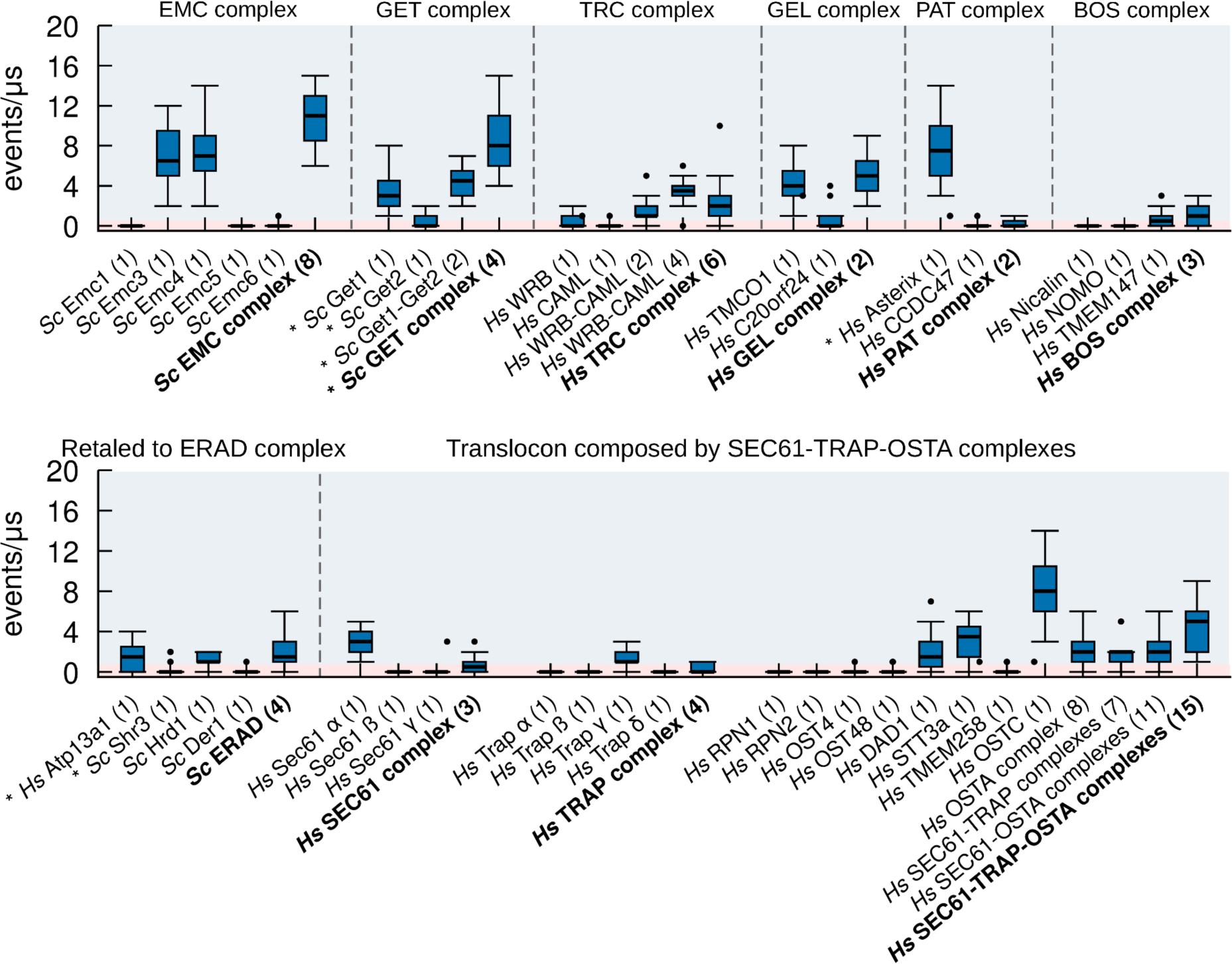
Extended version of insertase complexes in the endoplasmic reticulum. Lipid scrambling activity of individual components and selected complexes for the major ER insertases investigated. The blue shaded region represents the behavior for a scrambling-positive protein and the red area for a scrambling-negative protein. AlphaFold structures are denoted by the * symbol, oligomerization state is in parenthesis.

**Figure S7.**
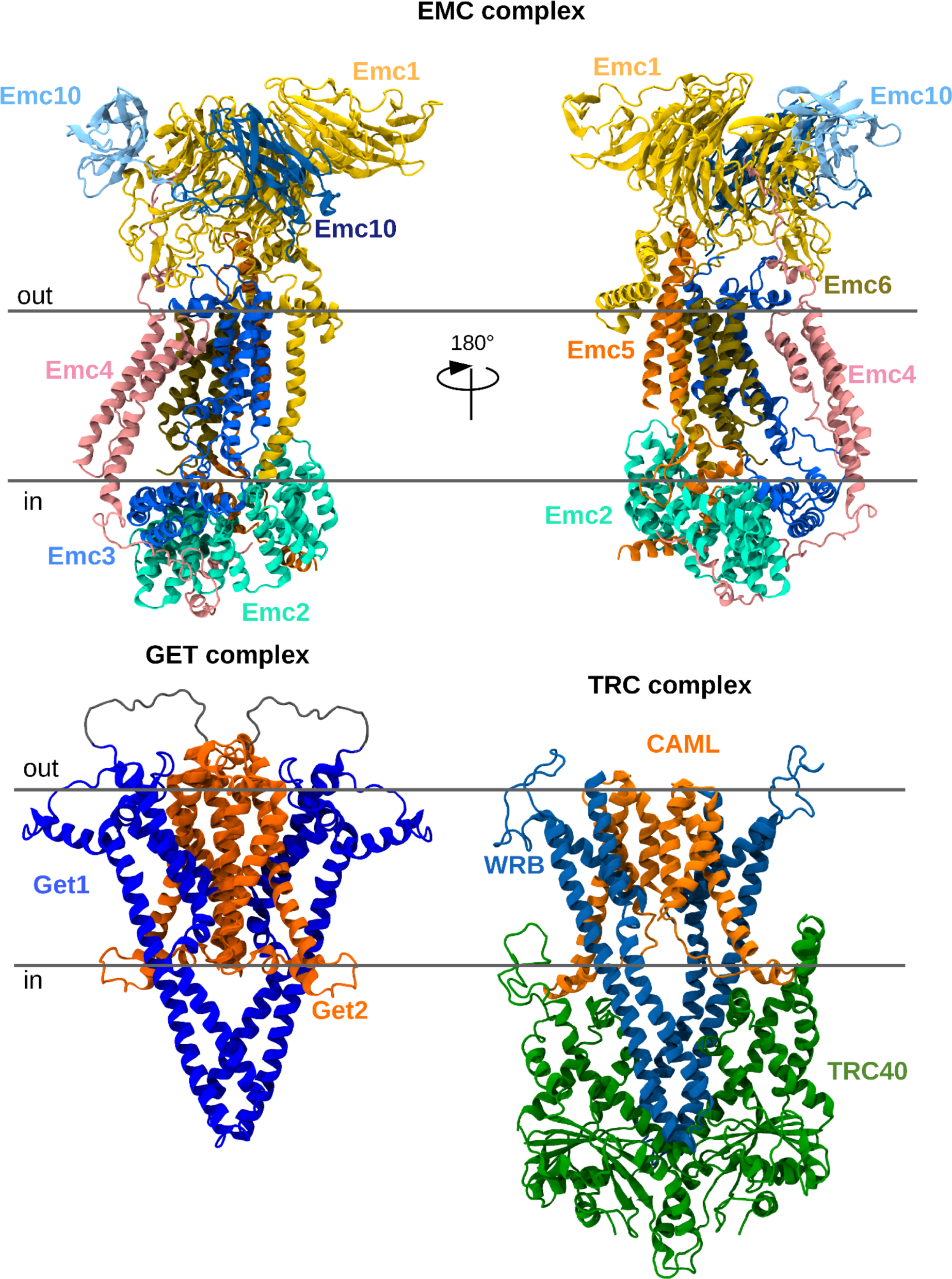
3D structures of ER complexes EMC, GET and TRC.

**Figure S8.**
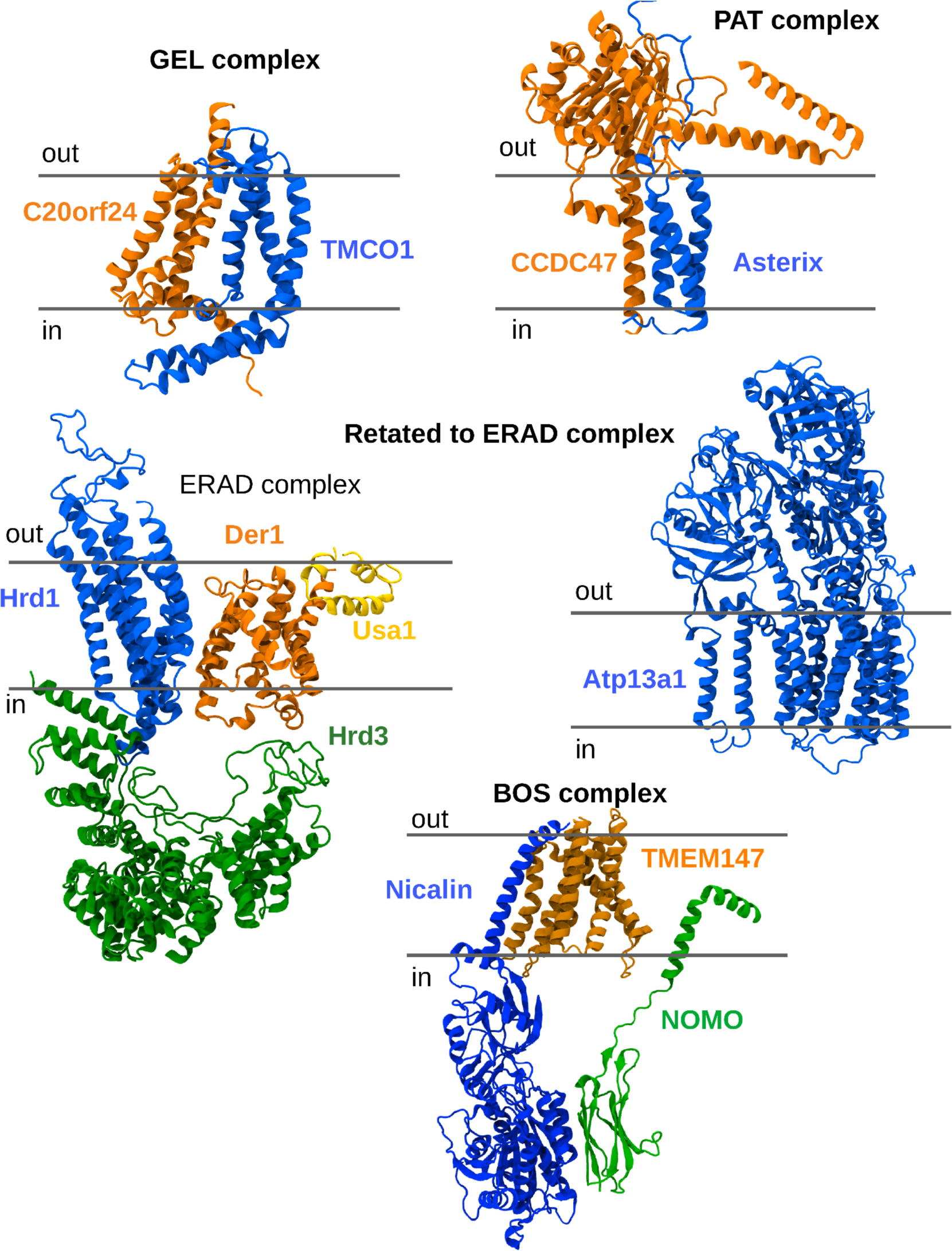
3D structures of ER complexes GEL, PAT, ERAD, Atp13a1 and BOS complex.

**Figure S9.**
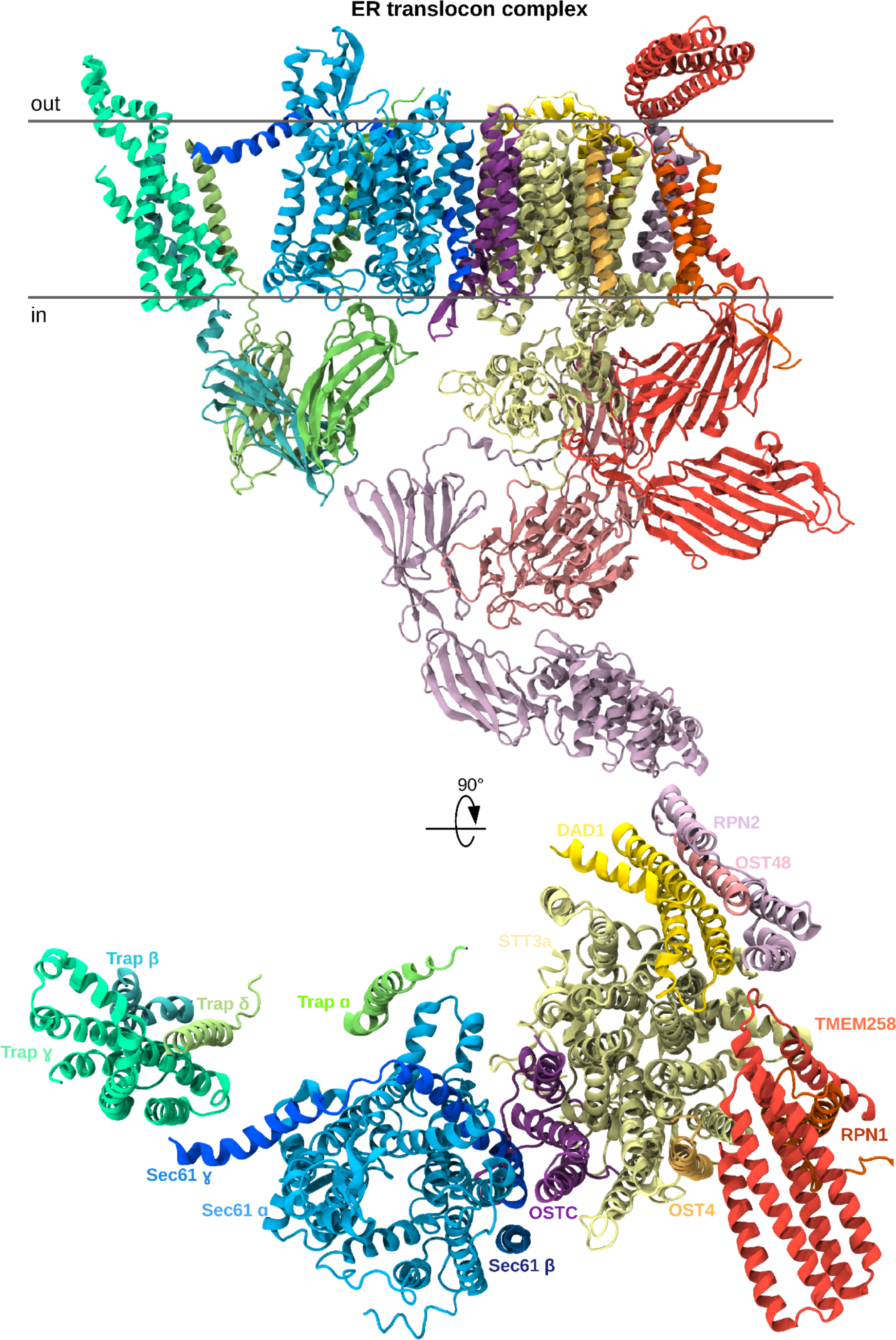
3D structure of *Hs*SEC61-TRAP-OSTA complex.

**Figure S10.**
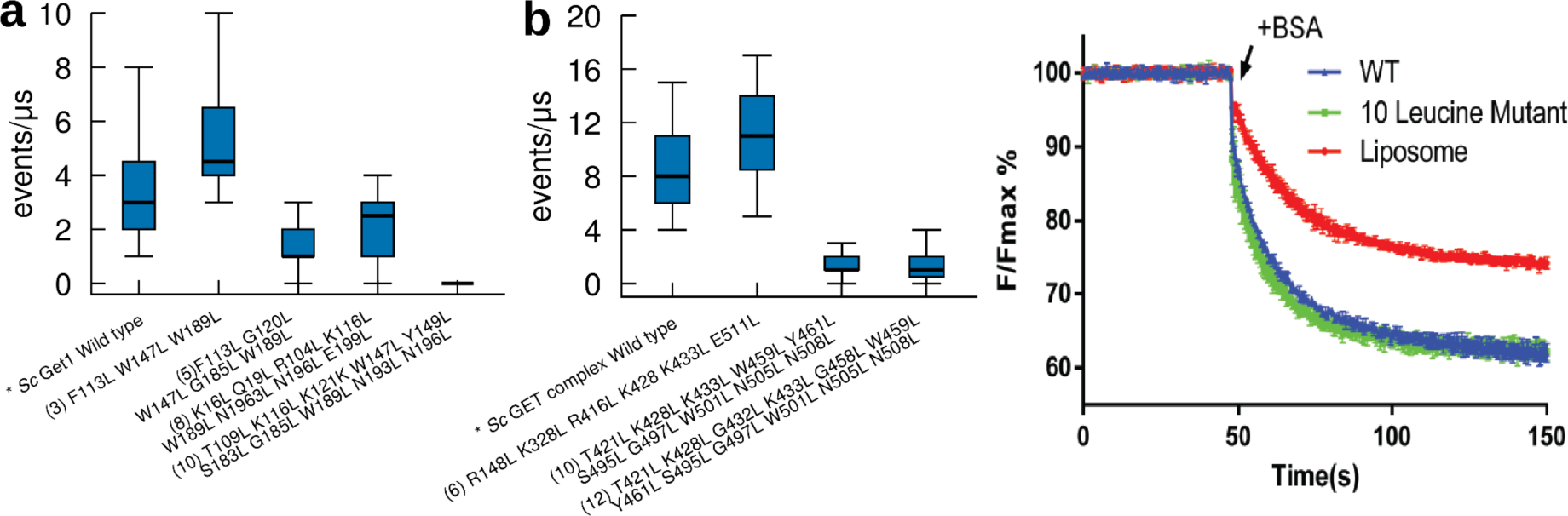
Mutational analysis of lipid scrambling in Get1 and GET complex. **a.** Scrambling activity calculations from CG-MD simulations of the wild type and various mutants of the *Sc*Get1 subunit. 10 mutations (10L) are required to completely ablate lipid scrambling activity in *Sc*Get1 **b.** Left. Scrambling activity calculations from CG-MD simulations of the wild type and various mutants of the *Sc*GET complex. Neither the same 10L mutations on Get1 and a new 12L mutations were able to ablate lipid scrambling activity in silico in the *Sc*GET complex. Right. Lipid scrambling activity of wt and L10 mutant GET. The scramblase assay for Get2-1sc mutant was carried out with FluoroMax+ spectrofluorometer (HORIBA). For each reaction, 50uL of the proteoliposomes were added to 1950uL of the reconstitution buffer. The sample was vigorously stirred and measured for fluorescence at 460/538 nm for 50-70 seconds to establish a stable baseline. 50uL of 1.5mg/mL fatty acid-free BSA was added, and fluorescence data were collected for another 200s. Both the WT complex and the mutant version with 10 mutations in the translocation channel (T421L/K428L/K433L/W459L/Y461L/S495L/G497L/W501L/N505L/N508L in Get1) scramble NBD-PC. **a-b**. The total number of mutations is in parentheses.

**Figure S11.**
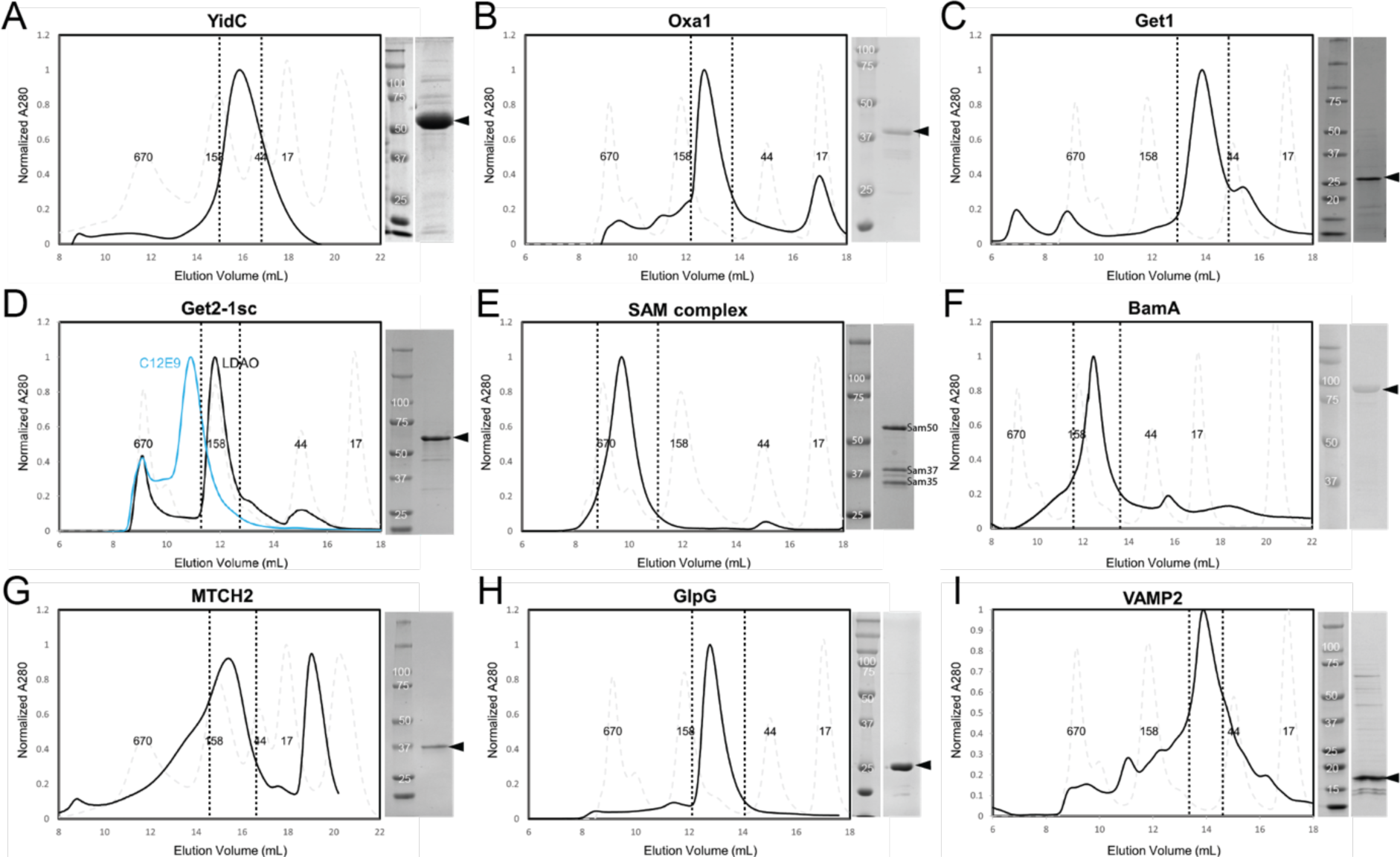
Size-exclusion chromatography analyses of proteins used in reconstitutions. Proteins for in vitro reconstitution were purified by size exclusion chromatography. Traces are shown (black) along with traces of the molecular weight standards (dotted grey); fractions used for reconstitutions are indicated (dotted black lines). Note that the elution volume of each protein reflects its size including detergent micelle. SDS-PAGE analysis of the purified protein is to the right of each chromatographic trace.

## References

Abraham, M.J., Murtola, T., Schulz, R., Páll, S., Smith, J.C., Hess, B., Lindahl, E., 2015. GROMACS: High performance molecular simulations through multi-level parallelism from laptops to supercomputers. SoftwareX 1–2, 19–25. 10.1016/j.softx.2015.06.001

Adlakha, J., Hong, Z., Li, P., Reinisch, K.M., 2022. Structural and biochemical insights into lipid transport by VPS13 proteins. J. Cell Biol. 221, e202202030. 10.1083/jcb.202202030

Anghel, S.A., McGilvray, P.T., Hegde, R.S., Keenan, R.J., 2017. Identification of Oxa1 Homologs Operating in the Eukaryotic Endoplasmic Reticulum. Cell Rep. 21, 3708–3716. 10.1016/j.celrep.2017.12.006

Antonicka, H., Lin, Z.-Y., Janer, A., Aaltonen, M.J., Weraarpachai, W., Gingras, A.-C., Shoubridge, E.A., 2020. A High-Density Human Mitochondrial Proximity Interaction Network. Cell Metab. 32, 479–497.e9. 10.1016/j.cmet.2020.07.017

Bai, L., You, Q., Feng, X., Kovach, A., Li, H., 2020. Structure of the ER membrane complex, a transmembrane-domain insertase. Nature 584, 475–478. 10.1038/s41586-020-2389-3

Brunner, J.D., Lim, N.K., Schenck, S., Duerst, A., Dutzler, R., 2014. X-ray structure of a calcium-activated TMEM16 lipid scramblase. Nature 516, 207–212. 10.1038/nature13984

Bushell, S.R., Pike, A.C.W., Falzone, M.E., Rorsman, N.J.G., Ta, C.M., Corey, R.A., Newport, T.D., Christianson, J.C., Scofano, L.F., Shintre, C.A., Tessitore, A., Chu, A., Wang, Q., Shrestha, L., Mukhopadhyay, S.M.M., Love, J.D., Burgess-Brown, N.A., Sitsapesan, R., Stansfeld, P.J., Huiskonen, J.T., Tammaro, P., Accardi, A., Carpenter, E.P., 2019. The structural basis of lipid scrambling and inactivation in the endoplasmic reticulum scramblase TMEM16K. Nat. Commun. 10, 3956. 10.1038/s41467-019-11753-1

Bussi, G., Donadio, D., Parrinello, M., 2007. Canonical sampling through velocity rescaling. J. Chem. Phys. 126, 014101. 10.1063/1.2408420

Chang, Q., Gummadi, S.N., Menon, A.K., 2004. Chemical Modification Identifies Two Populations of Glycerophospholipid Flippase in Rat Liver ER. Biochemistry 43, 10710–10718. 10.1021/bi049063a

Chua, G.-L., Tan, B.C., Loke, R.Y.J., He, M., Chin, C.-F., Wong, B.H., Kuk, A.C.Y., Ding, M., Wenk, M.R., Guan, L., Torta, F., Silver, D.L., 2023. Mfsd2a utilizes a flippase mechanism to mediate omega-3 fatty acid lysolipid transport. Proc. Natl. Acad. Sci. 120, e2215290120. 10.1073/pnas.2215290120

Evans, R., O’Neill, M., Pritzel, A., Antropova, N., Senior, A., Green, T., Žídek, A., Bates, R., Blackwell, S., Yim, J., Ronneberger, O., Bodenstein, S., Zielinski, M., Bridgland, A., Potapenko, A., Cowie, A., Tunyasuvunakool, K., Jain, R., Clancy, E., Kohli, P., Jumper, J., Hassabis, D., 2021. Protein complex prediction with AlphaFold-Multimer (preprint). Bioinformatics. 10.1101/2021.10.04.463034

Falzone, M.E., Feng, Z., Alvarenga, O.E., Pan, Y., Lee, B., Cheng, X., Fortea, E., Scheuring, S., Accardi, A., 2022. TMEM16 scramblases thin the membrane to enable lipid scrambling. Nat. Commun. 13, 2604. 10.1038/s41467-022-30300-z

Galazzo, L., Meier, G., Januliene, D., Parey, K., De Vecchis, D., Striednig, B., Hilbi, H., Schäfer, L.V., Kuprov, I., Moeller, A., Bordignon, E., Seeger, M.A., 2022. The ABC transporter MsbA adopts the wide inward-open conformation in *E. coli* cells. Sci. Adv. 8, eabn6845. 10.1126/sciadv.abn6845

Galietta, L.J.V., 2009. The TMEM16 protein family: a new class of chloride channels? Biophys. J. 97, 3047– 3053. 10.1016/j.bpj.2009.09.02

Gemmer, M., Chaillet, M.L., Van Loenhout, J., Cuevas Arenas, R., Vismpas, D., Gröllers-Mulderij, M., Koh, F.A., Albanese, P., Scheltema, R.A., Howes, S.C., Kotecha, A., Fedry, J., Förster, F., 2023. Visualization of translation and protein biogenesis at the ER membrane. Nature 614, 160–167. 10.1038/s41586-022-05638-5

Gentle, I., Gabriel, K., Beech, P., Waller, R., Lithgow, T., 2004. The Omp85 family of proteins is essential for outer membrane biogenesis in mitochondria and bacteria. J. Cell Biol. 164, 19–24. 10.1083/jcb.200310092

Gérard, S.F., Hall, B.S., Zaki, A.M., Corfield, K.A., Mayerhofer, P.U., Costa, C., Whelligan, D.K., Biggin, P.C., Simmonds, R.E., Higgins, M.K., 2020. Structure of the Inhibited State of the Sec Translocon. Mol. Cell 79, 406–415.e7. 10.1016/j.molcel.2020.06.013

Ghanbarpour, A., Valverde, D.P., Melia, T.J., Reinisch, K.M., 2021. A model for a partnership of lipid transfer proteins and scramblases in membrane expansion and organelle biogenesis. Proc. Natl. Acad. Sci. 118, e2101562118. 10.1073/pnas.2101562118

González Montoro, A., Auffarth, K., Hönscher, C., Bohnert, M., Becker, T., Warscheid, B., Reggiori, F., Van Der Laan, M., Fröhlich, F., Ungermann, C., 2018. Vps39 Interacts with Tom40 to Establish One of Two Functionally Distinct Vacuole-Mitochondria Contact Sites. Dev. Cell 45, 621–636.e7. 10.1016/j.devcel.2018.05.011

Guna, A., Stevens, T.A., Inglis, A.J., Replogle, J.M., Esantsi, T.K., Muthukumar, G., Shaffer, K.C.L., Wang, M.L., Pogson, A.N., Jones, J.J., Lomenick, B., Chou, T.-F., Weissman, J.S., Voorhees, R.M., 2022. MTCH2 is a mitochondrial outer membrane protein insertase. Science 378, 317–322. 10.1126/science.add1856

Hartzell, H.C., Yu, K., Xiao, Q., Chien, L.-T., Qu, Z., 2009. Anoctamin/TMEM16 family members are Ca2+-activated Cl-channels. J. Physiol. 587, 2127–2139. 10.1113/jphysiol.2008.163709

He, M., Kuk, A.C.Y., Ding, M., Chin, C.F., Galam, D.L.A., Nah, J.M., Tan, B.C., Yeo, H.L., Chua, G.L., Benke, P.I., Wenk, M.R., Ho, L., Torta, F., Silver, D.L., 2022. Spns1 is a lysophospholipid transporter mediating lysosomal phospholipid salvage. Proc. Natl. Acad. Sci. 119, e2210353119. 10.1073/pnas.2210353119

Holthuis, J.C.M., Jahn, H., Menon, A.K., Mizushima, N., 2022. An alliance between lipid transfer proteins and scramblases for membrane expansion. Fac. Rev. 11. 10.12703/r-01-0000015

Huang, D., Xu, B., Liu, L., Wu, L., Zhu, Y., Ghanbarpour, A., Wang, Y., Chen, F.-J., Lyu, J., Hu, Y., Kang, Y., Zhou, W., Wang, X., Ding, W., Li, X., Jiang, Z., Chen, J., Zhang, Xu, Zhou, H., Li, J.Z., Guo, C., Zheng, W., Zhang, Xiuqin, Li, P., Melia, T., Reinisch, K., Chen, X.-W., 2021. TMEM41B acts as an ER scramblase required for lipoprotein biogenesis and lipid homeostasis. Cell Metab. 33, 1655–1670.e8. 10.1016/j.cmet.2021.05.006

Itoh, Y., Andréll, J., Choi, A., Richter, U., Maiti, P., Best, R.B., Barrientos, A., Battersby, B.J., Amunts, A., 2021. Mechanism of membrane-tethered mitochondrial protein synthesis. Science 371, 846–849. 10.1126/science.abe0763

Itskanov, S., Wang, L., Junne, T., Sherriff, R., Xiao, L., Blanchard, N., Shi, W.Q., Forsyth, C., Hoepfner, D., Spiess, M., Park, E., 2023. A common mechanism of Sec61 translocon inhibition by small molecules. Nat. Chem. Biol. 10.1038/s41589-023-01337-y

Jahn, H., Bartoš, L., Dearden, G.I., Dittman, J.S., Holthuis, J.C.M., Vácha, R., Menon, A.K., 2022. Phospholipids are imported into mitochondria by VDAC, a dimeric beta barrel scramblase (preprint). Biochemistry. 10.1101/2022.10.17.512472

Jo, S., Kim, T., Iyer, V.G., Im, W., 2008. CHARMM-GUI: A web-based graphical user interface for CHARMM. J. Comput. Chem. 29, 1859–1865. 10.1002/jcc.20945

Jumper, J., Evans, R., Pritzel, A., Green, T., Figurnov, M., Ronneberger, O., Tunyasuvunakool, K., Bates, R., Žídek, A., Potapenko, A., Bridgland, A., Meyer, C., Kohl, S.A.A.,Ballard, A.J., Cowie, A., Romera-Paredes, B., Nikolov, S., Jain, R., Adler, J., Back, T., Petersen, S., Reiman, D., Clancy, E., Zielinski, M., Steinegger, M., Pacholska, M., Berghammer, T., Bodenstein, S., Silver, D., Vinyals, O., Senior, A.W., Kavukcuoglu, K., Kohli, P., Hassabis, D., 2021. Highly accurate protein structure prediction with AlphaFold. Nature 596, 583–589. 10.1038/s41586-021-03819-2

Koch, C., Räschle, M., Prescianotto-Baschong, C., Spang, A., Herrmann, J.M., 2023. The ER-SURF pathway uses ER-mitochondria contact sites for protein targeting to mitochondria (preprint). Biochemistry. 10.1101/2023.08.10.552816

Kohler, R., Boehringer, D., Greber, B., Bingel-Erlenmeyer, R., Collinson, I., Schaffitzel, C., Ban, N., 2009. YidC and Oxa1 Form Dimeric Insertion Pores on the Translating Ribosome. Mol. Cell 34, 344–353. 10.1016/j.molcel.2009.04.019

Kornmann, B., Currie, E., Collins, S.R., Schuldiner, M., Nunnari, J., Weissman, J.S., Walter, P., 2009. An ER-mitochondria tethering complex revealed by a synthetic biology screen. Science 325, 477–481. 10.1126/science.1175088

Kubelt, J., Menon, A.K., Müller, P., Herrmann, A., 2002. Transbilayer Movement of Fluorescent Phospholipid Analogues in the Cytoplasmic Membrane of *Escherichia coli*. Biochemistry 41, 5605– 5612. 10.1021/bi0118714

Kumazaki, K., Chiba, S., Takemoto, M., Furukawa, A., Nishiyama, K., Sugano, Y., Mori, T., Dohmae, N., Hirata, K., Nakada-Nakura, Y., Maturana, A.D., Tanaka, Y., Mori, H., Sugita, Y., Arisaka, F., Ito, K., Ishitani, R., Tsukazaki, T., Nureki, O., 2014. Structural basis of Sec-independent membrane protein insertion by YidC. Nature 509, 516–520. 10.1038/nature13167

Labbé, K., Mookerjee, S., Le Vasseur, M., Gibbs, E., Lerner, C., Nunnari, J., 2021. The modified mitochondrial outer membrane carrier MTCH2 links mitochondrial fusion to lipogenesis. J. Cell Biol. 220, e202103122. 10.1083/jcb.202103122

Lahiri, S., Chao, J.T., Tavassoli, S., Wong, A.K.O., Choudhary, V., Young, B.P., Loewen, C.J.R., Prinz, W.A., 2014. A Conserved Endoplasmic Reticulum Membrane Protein Complex (EMC) Facilitates Phospholipid Transfer from the ER to Mitochondria. PLoS Biol. 12, e1001969. 10.1371/journal.pbio.1001969

Li, Y.E., Wang, Y., Du, X., Zhang, T., Mak, H.Y., Hancock, S.E., McEwen, H., Pandzic, E., Whan, R.M., Aw, Y.C., Lukmantara, I.E., Yuan, Y., Dong, X., Don, A., Turner, N., Qi, S., Yang, H., 2021. TMEM41B and VMP1 are scramblases and regulate the distribution of cholesterol and phosphatidylserine. J. Cell Biol. 220, e202103105. 10.1083/jcb.202103105

Lindorff-Larsen, K., Piana, S., Palmo, K., Maragakis, P., Klepeis, J.L., Dror, R.O., Shaw, D.E., 2010. Improved side-chain torsion potentials for the Amber ff99SB protein force field: Improved Protein Side-Chain Potentials. Proteins Struct. Funct. Bioinforma. 78, 1950–1958. 10.1002/prot.22711

Liu, X., Salokas, K., Tamene, F., Jiu, Y., Weldatsadik, R.G., Öhman, T., Varjosalo, M., 2018. An AP-MS- and BioID-compatible MAC-tag enables comprehensive mapping of protein interactions and subcellular localizations. Nat. Commun. 9, 1188. 10.1038/s41467-018-03523-2

Maeda, S., Yamamoto, H., Kinch, L.N., Garza, C.M., Takahashi, S., Otomo, C., Grishin, N.V., Forli, S., Mizushima, N., Otomo, T., 2020. Structure, lipid scrambling activity and role in autophagosome formation of ATG9A. Nat. Struct. Mol. Biol. 27, 1194–1201. 10.1038/s41594-020-00520-2

Mannella, C.A., 1992. The “ins” and “outs” of mitochondrial membrane channels. Trends Biochem. Sci. 17, 315–320. 10.1016/0968-0004(92)90444-e

Matoba, K., Kotani, T., Tsutsumi, A., Tsuji, T., Mori, T., Noshiro, D., Sugita, Y., Nomura, N., Iwata, S., Ohsumi, Y., Fujimoto, T., Nakatogawa, H., Kikkawa, M., Noda, N.N., 2020. Atg9 is a lipid scramblase that mediates autophagosomal membrane expansion. Nat. Struct. Mol. Biol. 27, 1185– 1193. 10.1038/s41594-020-00518-w

McDowell, M.A., Heimes, M., Fiorentino, F., Mehmood, S., Farkas, Á., Coy-Vergara, J., Wu, D., Bolla, J.R., Schmid, V., Heinze, R., Wild, K., Flemming, D., Pfeffer, S., Schwappach, B., Robinson, C.V., Sinning, I., 2020. Structural Basis of Tail-Anchored Membrane Protein Biogenesis by the GET Insertase Complex. Mol. Cell 80, 72–86.e7. 10.1016/j.molcel.2020.08.012

McGilvray, P.T., Anghel, S.A., Sundaram, A., Zhong, F., Trnka, M.J., Fuller, J.R., Hu, H., Burlingame, A.L., Keenan, R.J., 2020. An ER translocon for multi-pass membrane protein biogenesis. eLife 9, e56889. 10.7554/eLife.56889

Meisinger, C., Rissler, M., Chacinska, A., Szklarz, L.K.S., Milenkovic, D., Kozjak, V., Schönfisch, B., Lohaus, C., Meyer, H.E., Yaffe, M.P., Guiard, B., Wiedemann, N., Pfanner, N., 2004. The mitochondrial morphology protein Mdm10 functions in assembly of the preprotein translocase of the outer membrane. Dev. Cell 7, 61–71. 10.1016/j.devcel.2004.06.003

Melia, T.J., Reinisch, K.M., 2022. A possible role for VPS13-family proteins in bulk lipid transfer, membrane expansion and organelle biogenesis. J. Cell Sci. 135, jcs259357. 10.1242/jcs.259357

Menon, A.K., Watkins, W.E., Hrafnsdóttir, S., 2000. Specific proteins are required to translocate phosphatidylcholine bidirectionally across the endoplasmic reticulum. Curr. Biol. 10, 241–252. 10.1016/S0960-9822(00)00356-0

Menon, I., Huber, T., Sanyal, S., Banerjee, S., Barré, P., Canis, S., Warren, J.D., Hwa, J., Sakmar, T.P., Menon, A.K., 2011. Opsin Is a Phospholipid Flippase. Curr. Biol. 21, 149–153. 10.1016/j.cub.2010.12.031

Mirdita, M., Schütze, K., Moriwaki, Y., Heo, L., Ovchinnikov, S., Steinegger, M., 2022. ColabFold: making protein folding accessible to all. Nat. Methods 19, 679–682. 10.1038/s41592-022-01488-1

Ni, D., Huang, Y., 2015. The Expression, Purification, and Structure Determination of BamA from E. coli. Methods Mol. Biol. Clifton NJ 1329, 169–178. 10.1007/978-1-4939-2871-2_13

Nosol, K., Romane, K., Irobalieva, R.N., Alam, A., Kowal, J., Fujita, N., Locher, K.P., 2020. Cryo-EM structures reveal distinct mechanisms of inhibition of the human multidrug transporter ABCB1. Proc. Natl. Acad. Sci. 117, 26245–26253. 10.1073/pnas.2010264117

Olsen, J.A., Alam, A., Kowal, J., Stieger, B., Locher, K.P., 2020. Structure of the human lipid exporter ABCB4 in a lipid environment. Nat. Struct. Mol. Biol. 27, 62–70. 10.1038/s41594-019-0354-3

Osawa, T., Kotani, T., Kawaoka, T., Hirata, E., Suzuki, K., Nakatogawa, H., Ohsumi, Y., Noda, N.N., 2019. Atg2 mediates direct lipid transfer between membranes for autophagosome formation. Nat. Struct. Mol. Biol. 26, 281–288. 10.1038/s41594-019-0203-4

Parrinello, M., Rahman, A., 1981. Polymorphic transitions in single crystals: A new molecular dynamics method. J. Appl. Phys. 52, 7182–7190. 10.1063/1.328693

Perez, C., Gerber, S., Boilevin, J., Bucher, M., Darbre, T., Aebi, M., Reymond, J.-L., Locher, K.P., 2015. Structure and mechanism of an active lipid-linked oligosaccharide flippase. Nature 524, 433–438. 10.1038/nature14953

Ploier, B., Menon, A.K., 2016. A Fluorescence-based Assay of Phospholipid Scramblase Activity. J. Vis. Exp. 54635. 10.3791/54635

Pomorski, T., Menon, A.K., 2006. Lipid flippases and their biological functions. Cell. Mol. Life Sci. 63, 2908– 2921. 10.1007/s00018-006-6167-7

Qi, L., Wang, Q., Guan, Z., Wu, Y., Shen, C., Hong, S., Cao, J., Zhang, X., Yan, C., Yin, P., 2021. Cryo-EM structure of the human mitochondrial translocase TIM22 complex. Cell Res. 31, 369–372. 10.1038/s41422-020-00400-w

Rapoport, T.A., Li, L., Park, E., 2017. Structural and Mechanistic Insights into Protein Translocation. Annu. Rev. Cell Dev. Biol. 33, 369–390. 10.1146/annurev-cellbio-100616-060439

Reinisch, K.M., Prinz, W.A., 2021. Mechanisms of nonvesicular lipid transport. J. Cell Biol. 220, e202012058. 10.1083/jcb.202012058

Ren, S., De Kok, N.A.W., Gu, Y., Yan, W., Sun, Q., Chen, Y., He, J., Tian, L., Andringa, R.L.H., Zhu, X., Tang, M., Qi, S., Xu, H., Ren, H., Fu, X., Minnaard, A.J., Yang, S., Zhang, W., Li, W., Wei, Y., Driessen, A.J.M., Cheng, W., 2020. Structural and Functional Insights into an Archaeal Lipid Synthase. Cell Rep. 33, 108294. 10.1016/j.celrep.2020.108294

Ruggles, K.V., Garbarino, J., Liu, Y., Moon, J., Schneider, K., Henneberry, A., Billheimer, J., Millar, J.S., Marchadier, D., Valasek, M.A., Joblin-Mills, A., Gulati, S., Munkacsi, A.B., Repa, J.J., Rader, D., Sturley, S.L., 2014. A Functional, Genome-wide Evaluation of Liposensitive Yeast Identifies the “RE2 Required for Viability” (ARV1) Gene Product as a Major Component of Eukaryotic Fatty Acid Resistance. J. Biol. Chem. 289, 4417–4431. 10.1074/jbc.M113.515197

Sakuragi, T., Kanai, R., Tsutsumi, A., Narita, H., Onishi, E., Nishino, K., Miyazaki, T., Baba, T., Kosako, H., Nakagawa, A., Kikkawa, M., Toyoshima, C., Nagata, S., 2021. The tertiary structure of the human Xkr8–Basigin complex that scrambles phospholipids at plasma membranes. Nat. Struct. Mol. Biol. 28, 825–834. 10.1038/s41594-021-00665-8

Sakuragi, T., Nagata, S., 2023. Regulation of phospholipid distribution in the lipid bilayer by flippases and scramblases. Nat. Rev. Mol. Cell Biol. 24, 576–596. 10.1038/s41580-023-00604-z

Serek, J., Bauer-Manz, G., Struhalla, G., van den Berg, L., Kiefer, D., Dalbey, R., Kuhn, A., 2004. Escherichia coli YidC is a membrane insertase for Sec-independent proteins. EMBO J. 23, 294–301. 10.1038/sj.emboj.7600063

Shao, S., 2023. Protein biosynthesis at the ER: finding the right accessories. Mol. Biol. Cell 34, pe1. 10.1091/mbc.E21-09-0451

Siggel, M., Bhaskara, R.M., Hummer, G., 2019. Phospholipid Scramblases Remodel the Shape of Asymmetric Membranes. J. Phys. Chem. Lett. 10, 6351–6354. 10.1021/acs.jpclett.9b02531

Sim, S.I., Chen, Y., Lynch, D.L., Gumbart, J.C., Park, E., 2023. Structural basis of mitochondrial protein import by the TIM23 complex. Nature. 10.1038/s41586-023-06239-6

Smalinskaitė, L., Kim, M.K., Lewis, A.J.O., Keenan, R.J., Hegde, R.S., 2022. Mechanism of an intramembrane chaperone for multipass membrane proteins. Nature 611, 161–166. 10.1038/s41586-022-05336-2

Souza, P.C.T., Alessandri, R., Barnoud, J., Thallmair, S., Faustino, I., Grünewald, F., Patmanidis, I., Abdizadeh, H., Bruininks, B.M.H., Wassenaar, T.A., Kroon, P.C., Melcr, J., Nieto, V., Corradi, V., Khan, H.M., Domański, J., Javanainen, M., Martinez-Seara, H., Reuter, N., Best, R.B., Vattulainen I., Monticelli, L., Periole, X., Tieleman, D.P., de Vries, A.H., Marrink, S.J., 2021. Martini 3: a general purpose force field for coarse-grained molecular dynamics. Nat. Methods 18, 382–388. 10.1038/s41592-021-01098-3

Straub, M.S., Alvadia, C., Sawicka, M., Dutzler, R., 2021. Cryo-EM structures of the caspase-activated protein XKR9 involved in apoptotic lipid scrambling. eLife 10, e69800. 10.7554/eLife.69800

Sundaram, A., Yamsek, M., Zhong, F., Hooda, Y., Hegde, R.S., Keenan, R.J., 2022. Substrate-driven assembly of a translocon for multipass membrane proteins. Nature 611, 167–172. 10.1038/s41586-022-05330-8

Sundberg, E., Slagter, J.G., Fridborg, I., Cleary, S.P., Robinson, C., Coupland, G., 1997. ALBINO3, an Arabidopsis nuclear gene essential for chloroplast differentiation, encodes a chloroplast protein that shows homology to proteins present in bacterial membranes and yeast mitochondria. Plant Cell 9, 717–730. 10.1105/tpc.9.5.717

Suzuki, J., Denning, D.P., Imanishi, E., Horvitz, H.R., Nagata, S., 2013. Xk-Related Protein 8 and CED-8 Promote Phosphatidylserine Exposure in Apoptotic Cells. Science 341, 403–406. 10.1126/science.1236758

Suzuki, J., Umeda, M., Sims, P.J., Nagata, S., 2010. Calcium-dependent phospholipid scrambling by TMEM16F. Nature 468, 834–838. 10.1038/nature09583

Tan, J.X., Finkel, T., 2022. A phosphoinositide signalling pathway mediates rapid lysosomal repair. Nature 609, 815–821. 10.1038/s41586-022-05164-4

Tang, Z., Takahashi, Y., He, H., Hattori, T., Chen, C., Liang, X., Chen, H., Young, M.M., Wang, H.-G., 2019. TOM40 Targets Atg2 to Mitochondria-Associated ER Membranes for Phagophore Expansion. Cell Rep. 28, 1744–1757.e5. 10.1016/j.celrep.2019.07.036

Tucker, K., Park, E., 2019. Cryo-EM structure of the mitochondrial protein-import channel TOM complex at near-atomic resolution. Nat. Struct. Mol. Biol. 26, 1158–1166. 10.1038/s41594-019-0339-2

Valverde, D.P., Yu, S., Boggavarapu, V., Kumar, N., Lees, J.A., Walz, T., Reinisch, K.M., Melia, T.J., 2019. ATG2 transports lipids to promote autophagosome biogenesis. J. Cell Biol. 218, 1787–1798. 10.1083/jcb.201811139

Vance, D.E., Choy, P.C., Farren, S.B., Lim, P.H., Schneider, W.J., 1977. Asymmetry of phospholipid biosynthesis. Nature 270, 268–269. 10.1038/270268a0

Vance, J.E., 2015. Phospholipid Synthesis and Transport in Mammalian Cells. Traffic 16, 1–18. 10.1111/tra.12230

Wang, H., Levental, K.R., Lorent, J.H., Revathi, A.A., Levental, I., 2023. Lipid scrambling facilitates membrane vesiculation through decreasing membrane stiffness. Biophys. J. 122, 22a–23a. 10.1016/j.bpj.2022.11.347

Wang, L., Hou, W.-T., Chen, L., Jiang, Y.-L., Xu, D., Sun, L., Zhou, C.-Z., Chen, Y., 2020. Cryo-EM structure of human bile salts exporter ABCB11. Cell Res. 30, 623–625. 10.1038/s41422-020-0302-0

Wu, X., Rapoport, T.A., 2021. Translocation of Proteins through a Distorted Lipid Bilayer. Trends Cell Biol. 31, 473–484. 10.1016/j.tcb.2021.01.002

Wu, X., Siggel, M., Ovchinnikov, S., Mi, W., Svetlov, V., Nudler, E., Liao, M., Hummer, G., Rapoport, T.A., 2020. Structural basis of ER-associated protein degradation mediated by the Hrd1 ubiquitin ligase complex. Science 368, eaaz2449. 10.1126/science.aaz2449

Zalisko, B.E., Chan, C., Denic, V., Rock, R.S., Keenan, R.J., 2017. Tail-Anchored Protein Insertion by a Single Get1/2 Heterodimer. Cell Rep. 20, 2287–2293. 10.1016/j.celrep.2017.08.035

Zhang, Yutong, Ou, X., Wang, X., Sun, D., Zhou, X., Wu, X., Li, Q., Li, L., 2021. Structure of the mitochondrial TIM22 complex from yeast. Cell Res. 31, 366–368. 10.1038/s41422-020-00399-0

Zhang, Yuxia, Watanabe, S., Tsutsumi, A., Kadokura, H., Kikkawa, M., Inaba, K., 2021. Cryo-EM analysis provides new mechanistic insight into ATP binding to Ca ^2+^-ATPase SERCA2b. EMBO J. 40, e108482. 10.15252/embj.2021108482

Zhou, Z., Torres, M., Sha, H., Halbrook, C.J., Van Den Bergh, F., Reinert, R.B., Yamada, T., Wang, S., Luo, Y., Hunter, A.H., Wang, C., Sanderson, T.H., Liu, M., Taylor, A., Sesaki, H., Lyssiotis, C.A., Wu, J., Kersten, S., Beard, D.A., Qi, L., 2020. Endoplasmic reticulum–associated degradation regulates mitochondrial dynamics in brown adipocytes. Science 368, 54–60. 10.1126/science.aay2494

